# LEA_4 motifs function alone and in conjunction with synergistic cosolutes to protect a labile enzyme during desiccation

**DOI:** 10.1101/2024.09.04.611296

**Authors:** Vincent Nicholson, Kenny Nguyen, Edith Gollub, Mary McCoy, Feng Yu, Alex S. Holehouse, Shahar Sukenik, Thomas C. Boothby

## Abstract

Organisms from all kingdoms of life depend on Late Embryogenesis Abundant (LEA) proteins to survive desiccation. LEA proteins are divided into broad families distinguished by the presence of family-specific motif sequences. The LEA_4 family, characterized by eleven-residue motifs, plays a crucial role in the desiccation tolerance of numerous species. However, the role of these motifs in the function of LEA_4 proteins is unclear, with some studies finding that they recapitulate the function of full-length LEA_4 proteins *in vivo*, and other studies finding the opposite result. In this study, we characterize the ability of LEA_4 motifs to protect a desiccation-sensitive enzyme, citrate synthase, from loss of function during desiccation. We show here that LEA_4 motifs not only prevent the loss of function of citrate synthase during desiccation, but also that they can do so more robustly via synergistically interactions with cosolutes. Our analysis further suggests that cosolutes induce synergy with LEA_4 motifs in a manner that correlates with transfer free energy (TFE). This research advances our understanding of LEA_4 proteins by demonstrating that during desiccation their motifs can protect specific clients to varying degrees and that their protective capacity is modulated by their chemical environment. Our findings extend beyond the realm of desiccation tolerance, offering insights into the interplay between IDPs and cosolutes. By investigating the function of LEA_4 motifs, we highlight broader strategies for understanding protein stability and function.

## Introduction

While water is essential for life’s active processes, many organisms can persist for years, or even decades, in a desiccated (meaning dried) ametabolic state known as anhydrobiosis (“life without water”) [1]. How anhydrobiotic organisms tolerate desiccation, is an enduring paradox for biologists with broad implications for biotechnology/agriculture/etc.

A strategy often employed by organisms to survive desiccation is the enrichment of protective cosolutes, such as trehalose, glycine betaine, and glycerol, to a significant fraction (>10%) of the organism’s dry mass [2–5]. Cosolute mediators of desiccation tolerance protect cells and their labile components through a variety of mechanisms [6–8]. In several instances, the enrichment of cosolutes in anhydrobiotic organisms has been shown to be necessary and sufficient for conferring desiccation tolerance. For example, trehalose accumulation in stationary phase yeast is required for the acquisition of desiccation tolerance, while the exogenous introduction of this sugar makes normally desiccation-sensitive log-phase yeast tolerant to desiccation [9,10]. In addition, while some cosolute mediators of desiccation tolerance are widespread among different species of anhydrobiotic organisms, others are more limited in their taxonomic distribution [10,11]. For example, some desiccation-tolerant plants accumulate high levels of sucrose, which animals do not produce [6,8,12].

In addition to the enrichment of cosolutes, a more recent paradigm in the desiccation tolerance field is the accumulation of high levels of intrinsically disordered proteins (IDPs) [13–15]. While several families of desiccation-related IDPs have been identified [16–18], Late Embryogenesis Abundant (LEA) proteins are one of the most widely studied and characterized [19,20]. First identified as a mediator of abiotic stress in cotton seeds, LEA proteins have since been discovered across the kingdoms of life [16,21,22]. Moreover, previous work has demonstrated that cosolutes and desiccation-protective LEAs can act synergistically with one another, whereby their combined efficacy is greater than the sum of their individual contributions [23–25].

LEA proteins themselves are classified into different families based on the presence of conserved motifs. One family of LEA proteins, known as LEA_4 proteins, is characterized by the presence of 11mer motifs and are upregulated in desiccation-tolerant organisms during desiccation stress [16,26,27]. LEA_4 proteins are commonly cited for their ability to protect both labile proteins and membranes during desiccation [27]. These LEA proteins undergo a disorder-to-helix transition during desiccation [28,29], which is thought to be important for their protective function [28–31]. While the protective mechanism(s) employed by LEA_4 proteins are not fully elucidated, one working hypothesis is that they prevent protein aggregation through a process known as molecular shielding, in which protective proteins physically block interactions between aggregation prone clients [15].

The LEA_4 11mer motif has been proposed to be necessary and sufficient to recapitulate the behavior of full-length LEA_4 proteins [16,22]. For example, previous work found that replacing full-length LEA_4 proteins with just the LEA_4 motif is sufficient for desiccation tolerance in *C. elegans* [22]. In addition, LEA motifs from other families are capable of undergoing conserved structural transitions during desiccation [32]. With this in mind, an emerging model has suggested that LEA_4 motifs are the functional modules for desiccation protection within LEA_4 proteins [16,22].

Despite evidence that motifs from other LEA families recapitulate the functions and behaviors of full-length LEA proteins from the same family [22], for LEA_4 proteins and their motifs this may not be true. Recent work found that while full-length LEA_4 proteins robustly protect the desiccation-sensitive enzyme lactate dehydrogenase (LDH), LEA_4 motifs do not [25]. Moreover, LEA_4 motifs did not function synergistically with cosolutes, despite synergy being observed with many different full-length desiccation-related IDPs, including LEA_4 proteins [23–25,28,33]. One possible explanation for this apparent incongruity is that LEA_4 motifs may be sufficient to protect only a subset of clients stabilized by full-length LEA_4 proteins during desiccation. If this were the case, we may expect different desiccation-sensitive clients to be differentially protected by different LEA_4 motifs. However, prior studies into LEA_4 motifs’ *in vitro* protection have been limited to only LDH [25].

Here, we test the ability of LEA_4 motifs to protect another desiccation-sensitive enzyme, citrate synthase (CS), during desiccation. We find that in contrast to studies on LDH, four of seven LEA_4 motifs tested here conferred protection to CS above 50%. Additionally, we probe whether synergy with cosolutes is observed for LEA_4 motifs in protection of CS during desiccation. Again, in contrast to the previous observation that cosolutes do not enhance LEA_4 motif protection of LDH, we find that LEA_4 protection of CS is enhanced synergistically by several cosolutes. Finally, our results indicate that synergy correlates with the nature of the interaction between the cosolute and the motif, as measured by transfer free energy (TFE). Taken together, our work highlights the specificity of client:protectant interactions desiccation. Combined with previous findings, our work demonstrates that while full-length LEA_4 proteins protect a broad range of client enzymes, their motifs protect only a subset of these.

## Results

### LEA_4 motifs prevent citrate synthase (CS) loss of function during desiccation

While LEA_4 motifs do not protect the enzyme LDH during desiccation [25], to test whether they might protect other clients under these conditions, we selected seven model LEA_4 motifs derived from desiccation-tolerant organisms spanning different biological kingdoms (Table 1) and assessed their ability to preserve CS function during desiccation.

**Table 1:**
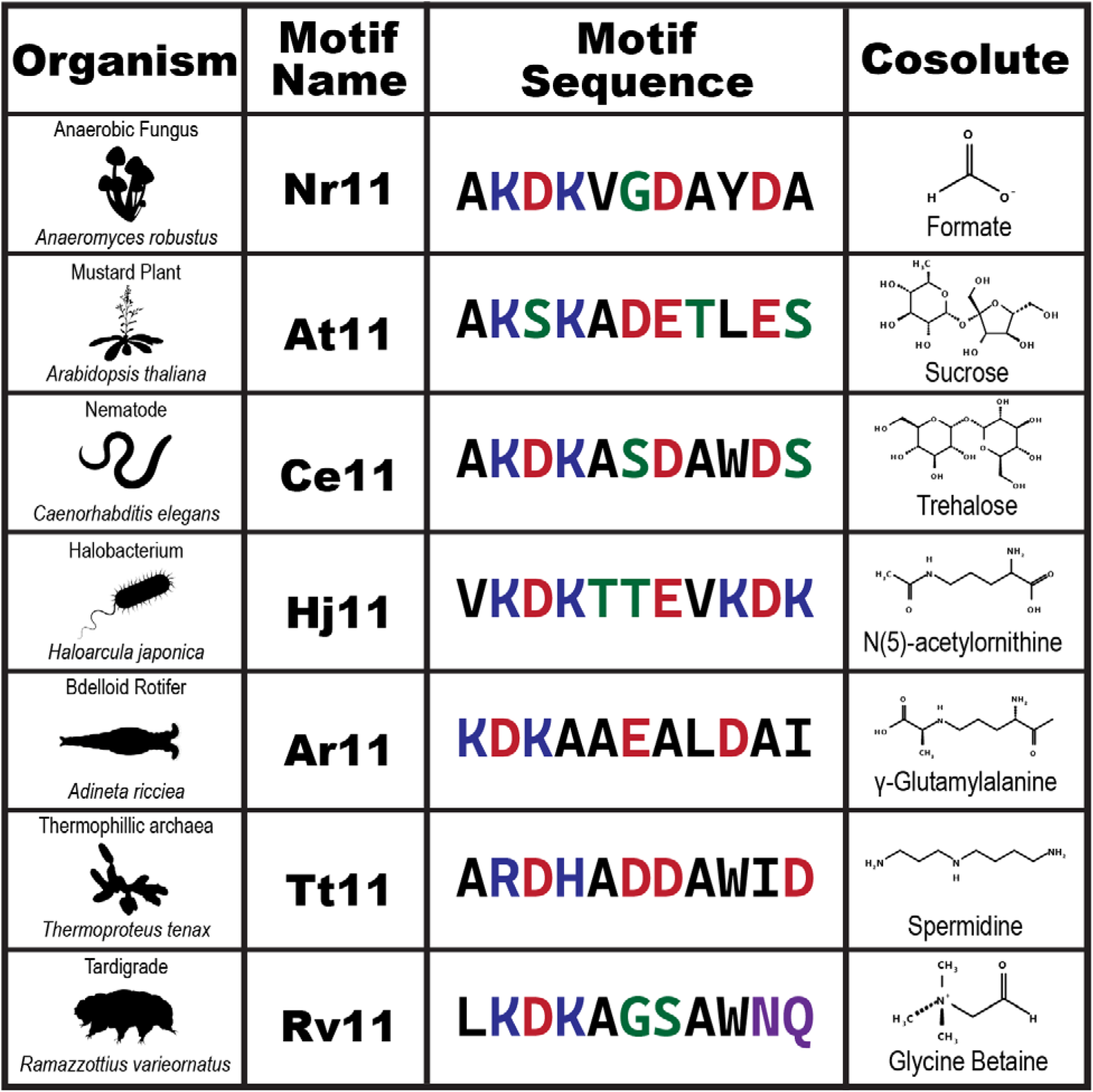
Representative cosolutes and LEA_4 motifs from seven organisms. Seven desiccation-tolerant organisms are shown alongside one of their LEA_4 motifs and one drying-enriched cosolute.

| Organism | Motif Name | Motif Sequence | Cosolute |
| --- | --- | --- | --- |
| Anaerobic Fungus<br>*Anaeromyces robustus* | Nr11 | AKDKVGDAYDA | Formate |
| Mustard Plant<br>*Arabidopsis thaliana* | At11 | AKSKADETLES | Sucrose |
| Nematode<br>*Caenorhabditis elegans* | Ce11 | AKDKASDAWDS | Trehalose |
| Halobacterium<br>*Haloarcula japonica* | Hj11 | VKDKTTEVKDK | N(5)-acetylornithine |
| Bdelloid Rotifer<br>*Adineta ricciae* | Ar11 | KDKAAEALDAI | γ-Glutamylalanine |
| Thermophilic archaea<br>*Thermoproteus tenax* | Tt11 | ARDHADDAWID | Spermidine |
| Tardigrade<br>*Ramazzottius varieornatus* | Rv11 | LKDKAGSAWNQ | Glycine Betaine |

Citrate synthase was incubated with each LEA_4 motif at several concentrations and then subjected to six cycles of desiccation and rehydration, which has previously been shown to be sufficient for complete loss of CS function [34] (Fig. 1). Four of the seven LEA_4 motifs tested preserved 50% or more of the enzymatic function of CS during desiccation (Nr11, Ce11, Tt11, Rv11; Fig. 1a-g). Some LEA_4 motifs exhibited a concentration-dependent increase in protection (Ce11; Fig. 1e), whereas others had an optimal concentration, resulting in non-monotonic protection (Nr11, At11, Tt11, Rv11; Fig. 1a,b,f,g). Two of the seven motifs displayed minimal protection to CS at the concentrations tested (Hj11, Ar11; Fig. 1 d,e).

**Figure 1:**
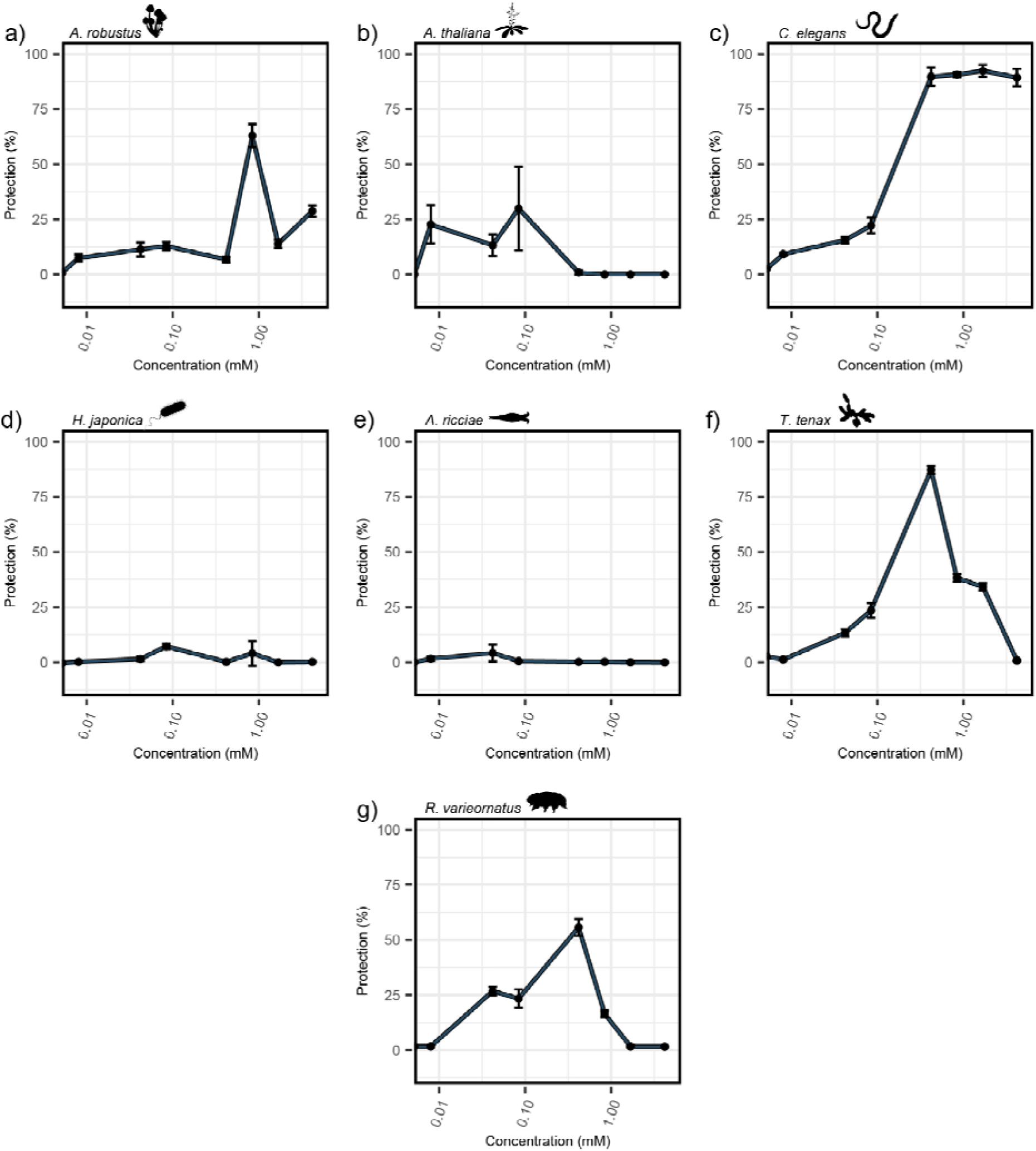
LEA_4 motifs are sufficient to protect citrate synthase from aggregation *in vitro*. The ability of LEA_4 motifs at various concentrations to protect 20 µM citrate synthase from si rounds of desiccation and rehydration. a) Nr11. b) At11. c) Ce11. d) Hj11. e) Ar11. f) Tt11. g) Rv11.

These results show that some LEA_4 motifs do provide robust protection to CS. These results also demonstrate that the degree of protection, as well as the optimal concentration of motif peptides, is sequence-dependent despite their homology.

### The sequence features of LEA_4 motifs do not correlate with protection

The results above (Fig. 1) demonstrate that CS protection varied depending on the LEA_4 motif sequence. Motivated by these observations, we next characterized sequence features of our seven LEA_4 motifs. Given that LEA_4 motifs are essentially simple intrinsically disordered proteins (IDPs), we wondered if principles developed in the context of IDPs would enable us to make sense of the sequence dependence shown in Fig. 1 (Fig. S1 & S2). Several sequence features are important to the function of intrinsically disordered proteins including charge patterning, hydropathy, and net charge [35–37].

The distribution of charges (kappa) in our seven motifs varies significantly, with kappa values ranging from 0 (well mixed) to 0.5 (highly segregated) (Fig. S1a). There is also significant variance in the average Kyte-Doolittle hydropathy; our LEA_4 motifs range from a near-neutral hydropathy (Ar11) to extremely hydrophilic (Hj11) (Fig. S1b). Finally, a Das-Pappu phase diagram of our motifs reveals diversity in the composition of their charged residues (Fig. S1c). The motifs vary in their fraction of charged residues from 27% to 63%. Two of the motifs have a net positive charge at pH = 7.0 (Hj11 and Rv11), while the rest are negatively charged (Fig. S1c).

Previous literature has highlighted the importance of these sequence features to the ensemble features, and sometimes the function, of IDPs [38,39]. Despite this, we have found no correlation between individual sequence features and the protective capacity of LEA_4 motifs for CS (Fig. S1d-h). This indicates that protection may arise from a LEA_4 motif’s structure and protein:protein or protein:solvent interactions as opposed to its linear sequence features.

### Desiccation-enriched cosolutes prevent citrate synthase (CS) loss of function

To test whether LEA_4 motifs can synergize with cosolutes to protect CS, we first tested the ability of cosolutes alone to preserve CS activity during desiccation. Using metabolomics data, we identified a desiccation-enriched cosolute from each of our seven organisms (Table 1) [4,24,40–43]. The resulting list included cosolutes already heavily implicated in stress tolerance, such as trehalose, sucrose, betaine, and spermidine [24,44]. It also included cosolutes outside the scope of normal stress tolerance literature, such as formate, N(5)-acetylornithine, and γ-glutamylalanine. In addition, several of these cosolutes are not specific to a single organism from Table 1. For example, trehalose is found in a wide range of desiccation-sensitive and - tolerant organisms [6].

We next tested the ability of cosolutes to protect CS at molar ratios ranging from 10:1 to 2000:1 of cosolute:CS. Like LEA_4 motifs, these seven cosolutes varied in their protective capacity (Fig. S3). Unlike in LEA_4 motifs, however, we see fewer non-monotonic trends in protection. The known desiccation tolerance mediators trehalose and sucrose protected CS in a concentration-dependent manner. Several other cosolutes provided robust protection at all concentrations examined, while others provided no protection at any concentration examined (Fig. S3). These results, combined with our motif protection data, provide baseline protection capacities for LEA_4 motifs and cosolutes alone - both of which are essential values for the synergy studies that we next carried out.

### LEA_4 motifs exhibit synergy with cosolutes

Given that LEA_4 motifs can protect CS during desiccation, we wondered if they might also function synergistically with co-enriched cosolutes.

The protective capacity of each protein and cosolute was first tested individually and then as a mixture (see example plot in Fig. S4a). Functional synergy occurs when a mixture protects significantly better than the sum of its individual parts (Fig. S4a). Functional antagonism occurs when a mixture protects significantly worse than the sum of its parts (Fig. S4a). This analysis was performed for each motif-cosolute combination, with the raw data available in Supplemental Figure S4 (Fig. S4b-l). From this data, we derived a ‘synergy index,’ which is calculated as the fractional difference between the mixture’s protection and the summed protection of the motif and cosolute alone (see *Methods*). Under this scheme, a synergy index of zero indicates protection of the mixture is merely additive, less than zero indicates functional antagonism, and greater than zero indicates functional synergy. This interpretation of the data can be flawed when the additive protection of both protein and cosolute is near 0% or 100%. Thus, concentrations for our synergy experiments have been selected specifically to avoid this issue, giving protection significantly greater than 0% but far less than 100% (Fig. S4).

A heatmap of the synergy index of each LEA_4 motif and cosolute combination reveals that protection of CS by LEA_4 motifs is heavily influenced by the inclusion of cosolutes (Fig. 2a). This influence was sometimes synergistic and other times antagonistic (Fig. 2a). Some cosolutes, such as formate and trehalose, resulted in synergistic interactions with all LEA_4 motifs. Betaine, on the other hand, was antagonistic in every combination examined. Other cosolutes, such as γ-glutamylalanine, sucrose, and spermidine, worked synergistically with some motifs and antagonistically with others (Fig. 2a). A heatmap of the protective capacity showed a similar result in which some cosolutes, like trehalose, tended to result in higher-protection mixtures, while others, like betaine, tended to be unprotective (Fig. 2b).

**Figure 2.**
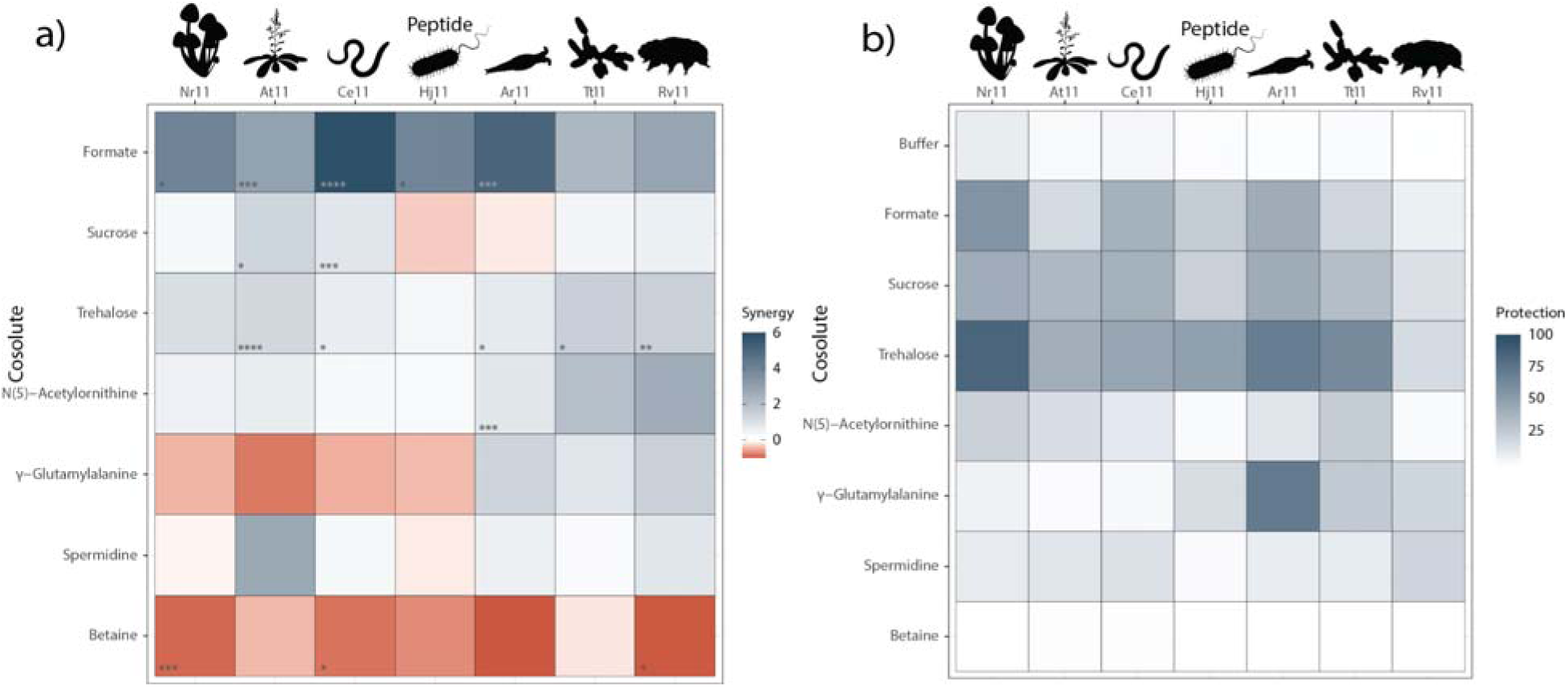
LEA_4 motifs are sufficient to protect against protein aggregation and exhibit robust synergy with specific cosolutes. **a)** Heatmap of synergy between 11-mer LEA_4 motifs and various cosolutes. Asterisks represent the statistical significance of synergy (see Fig. S4). Boxes without asterisks are not significantly different from additive protection. **b)** Heatmap of protection between 11-mer LEA_4 motifs and various cosolutes.

Because LEA_4 motifs are often found serially repeated in full-length LEA_4 proteins [45], we assessed the degree to which peptide motif repeat number had an effect on synergy, using the *A. thaliana* and *C. elegans* motifs as models to generate peptides with 2X (22mer) and 4X (44mer) repeats (Fig. S5a-d). Assessing synergy between 2X and 4X repeat peptides and cosolutes revealed that neither synergy nor protection change linearly with motif repeat number in the majority of cases (Fig. S5a-d). Instead, we often observed non-monotonic relationships for synergy, protection, or both (Fig. S5a-h).

Taken together, these results demonstrate that the protective capacity of LEA_4 motifs for CS during desiccation is heavily and differentially influenced by diverse cosolutes. This is in contrast to previous work showing that LEA_4 motifs offer minimal protection to LDH, and that protection of LDH by LEA_4 motifs is not enhanced by the addition of cosolutes that synergize with full-length LEA_4 proteins [25].

### Cosolutes effects on the global dimensions of LEA_4 motifs do not correlate with synergy

After observing synergy between LEA_4 motifs and some cosolutes in the protection of CS during desiccation, we became curious about the mechanisms underlying this synergy. Short peptides are inherently sensitive to their solution environment [46,47]. While they do not exist in a fixed three-dimensional structure, their environment can influence their conformational ensemble [48]. We, therefore, wondered if cosolute-induced synergy with LEA_4 motifs might be linked to a change in their ensemble.

The molecular shielding theory suggests that IDPs can shield sensitive biomolecules from aggregation by taking up space and slowing or preventing aggregation-prone proteins from interacting [15]. From this, one might reason that an IDP’s global dimensions will impact its protective capacity. For example, IDPs with more expanded radii of gyration might serve as better molecular shields by binding a larger protein surface area and preventing more interactions on average. Thus, a metabolite’s impact on the global dimensions of a LEA_4 motif may mechanistically underlie its synergistic interactions.

To test this, we assessed the global dimensions of LEA_4 motifs in the presence of different cosolutes using SAXS to determine the influence of these cosolutes on a motif’s radius of gyration (R_g_) [49–52]. These experiments were performed in high concentrations (0.438 M) of each cosolute, which approximate the crowded conditions experienced by LEA_4 proteins during the intermediate stages of desiccation. Some combinations of LEA_4 motifs and cosolutes resulted in low-quality scattering profiles for which an R_g_ could not be determined, and thus these scattering profiles were excluded from all analysis (see *Methods*). In the remaining data, we observed that the global dimensions of LEA_4 motifs were significantly affected by the solution environment (Fig. 3a). Notably, the vast majority of LEA_4 motifs became more compact when exposed to a cosolute. However, contrary to our assumption, we found no statistically significant correlation between the change in global dimensions (ΔR_g_) and functional synergy for each peptide-cosolute combination (Fig. 3b). We see a slightly better, but still weak correlation between ΔR_g_ and basal protection (Fig. 3c). Importantly, the changes in Rg observed here are small and near the limits of resolution for SAXS. Overall, our data does not conclusively suggest a relationship between synergistic protection of CS and the change in the radius of gyration of LEA_4 motifs.

**Figure 3.**
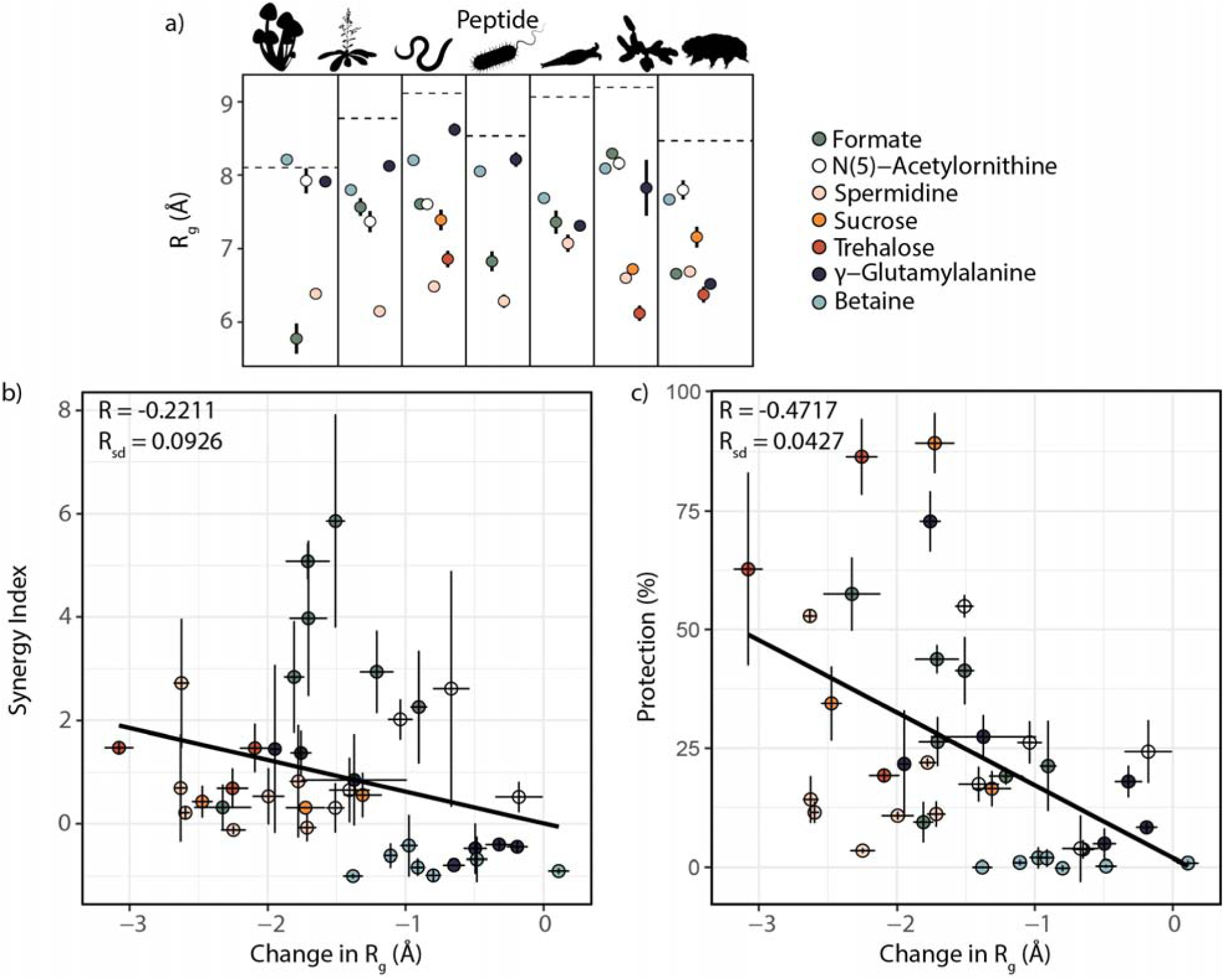
The influence of cosolutes on the global ensemble of LEA_4 motifs. **a)** The radius of gyration of LEA_4 motifs in the presence of 0.438 M of various cosolute. The dashed line represents the radius of gyration of the motif in a simple biological buffer. Black lines represent uncertainty in the radius of gyration (R_g_), as reported by BioXTAS RAW. **b)** Correlation between the change in radius of gyration and synergy index of peptide-cosolute mixtures. **c)** Correlation between the change in radius of gyration and percent protection of peptide-cosolute mixtures. For all correlations, R (correlation coefficient) was generated using a Pearson’s Correlation in R 4.3.0. R_sd_ (the standard deviation of the correlation coefficient) was generated using a Monte Carlo error simulation approach for correlations (see *Methods*).

### Cosolutes do not measurably affect the local ensemble of LEA_4 motifs in the hydrated state

Many LEA_4 motifs have been shown to undergo a coil-to-helix transition upon desiccation, which is thought to increase or confer protective function [16,28,30]. We therefore wondered if synergy observed in our CS assay could be caused by a change in the local secondary structure of LEA_4 motifs, whereby cosolutes pre-organize LEA_4 motifs with enhanced basal helicity. Importantly, these changes may not be observed using SAXS.

To test this, we performed circular dichroism (CD) spectroscopy on LEA_4 motifs in buffer and in the presence of 100 mM of trehalose, sucrose, and betaine. Qualitatively, these cosolutes appeared to elicit little to no change in the secondary structure of LEA_4 motifs (Fig. 4). To see if subtle differences could be detected in these spectra, we used two CD deconvolution tools, BeStSel and DichroWeb [53–56]. Both BeStSel (Fig. 4) and DichroWeb (Fig. S6) reveal no consistent change in the secondary structure of LEA_4 motifs in the presence of these cosolutes. Furthermore, BeStSel and DichroWeb disagree on the structural composition of the motifs in a plain buffer, as well as the extent to which structure changes upon the addition of cosolutes.

**Figure 4.**
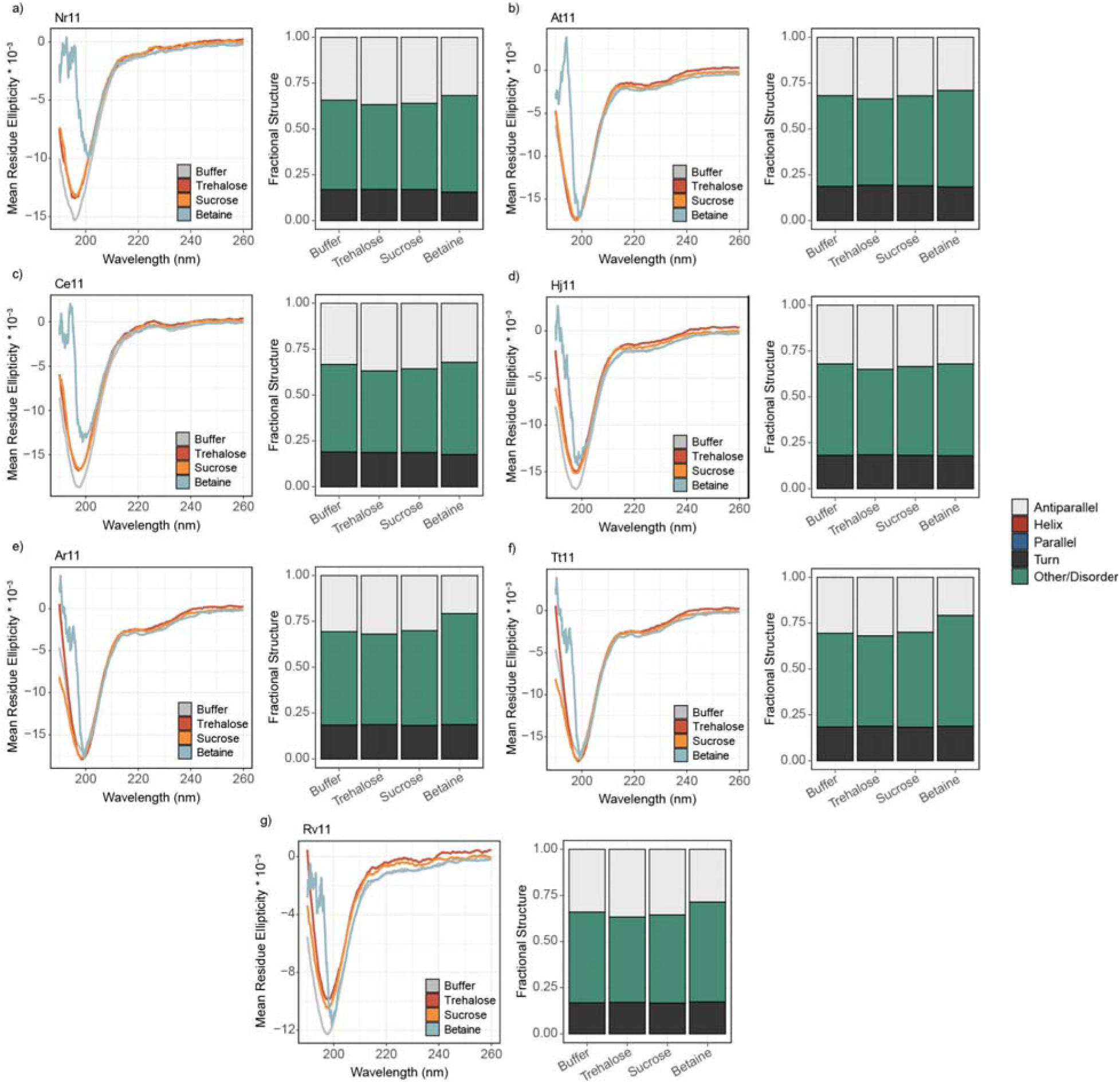
The influence of cosolutes on the local ensemble of LEA_4 motifs. CD Spectra for each LEA_4 11-mer motif in molar residue ellipticity (MRE) × 10^−3^ (left) and deconvoluted fractional secondary structure from BeStSel (right). **a)** Nr11, **b)** At11, **c)** Ce11, **d)** Hj11, **e)** Ar11, **f)** Tt11, **g)** Rv11.

Taken together, these results demonstrate no clear relationship between the presence of different cosolutes and specific changes in the structure of LEA_4 motifs. At the concentrations tested, the secondary structure of LEA_4 motifs was insensitive to the presence of cosolutes, implying that secondary structure is not a major mechanism through which cosolutes modulate LEA_4 motif function. Therefore, from these data we surmise that changes in secondary structure do not underlie the synergy observed between cosolutes and LEA_4 motifs in protecting CS during desiccation.

### Transfer free energy as a predictor of synergy

Given that our analysis thus far lacked a clear relationship between structure and protective function, we wondered if chemical interactions might explain the observed interplay between sequence, cosolute, and protection. Previous work demonstrated that transfer free energies (TFEs) can predict, and explain the mechanism underlying functional synergy between full-length LEA proteins and cosolutes [25]. TFEs measure the change in free energy of a molecule as it is transferred from water into a solution of a given cosolute at a standard state (typically 1 M) [57–59]. We used amino acid TFEs to calculate the repulsiveness or attractiveness of each cosolute to our LEA_4 motifs *ΔG_tr_* using the following equation:

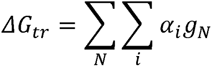

Here, *i* is a numerical index for all instances of the chemical group (defined as one of 19 amino-acid side chains and the peptide backbone), *α* is the relative solvent accessibility of the chemical group as a fraction of 1 (1 being completely solvent accessible and 0 being completely buried), *N* is the chemical group, and *g* is the experimental value of the transfer free energy for that chemical group [25,60,61]. For the purposes of this study, we assume *α* = 1 for all residues based on the CD-resolved disordered nature of LEA_4 motifs.

Of the seven cosolutes used in this study, only three (trehalose, sucrose, and betaine) have experimental transfer free energy data publicly available [57,62]. Calculating the *ΔG_tr_* of LEA_4 motifs with each of these three cosolutes reveals that they have a diverse range of values (Fig. 5a). In all cases, the *ΔG_tr_* of a LEA_4 motif with betaine is negative, indicating that protein:solvent interactions are stabilized relative to protein:protein interactions. The *ΔG_tr_* of LEA_4 motifs with trehalose and sucrose is positive, indicating that protein:protein interactions are stabilized relative to protein:solvent interactions (Fig. 5a). Simply put, these predictions suggest that betaine acts as an excellent solvent for LEA_4 motifs, potentially disrupting protein:protein interactions. Conversely, sucrose and trehalose have repulsive interactions with LEA_4 motifs, increasing the favorability of protein:protein interactions that would hide away some of their surface area. Such a protein:protein interactions could be intra-protein, inter-protein homotypic (motif:motif), or inter-protein heterotypic (motif:CS).

**Figure 5.**
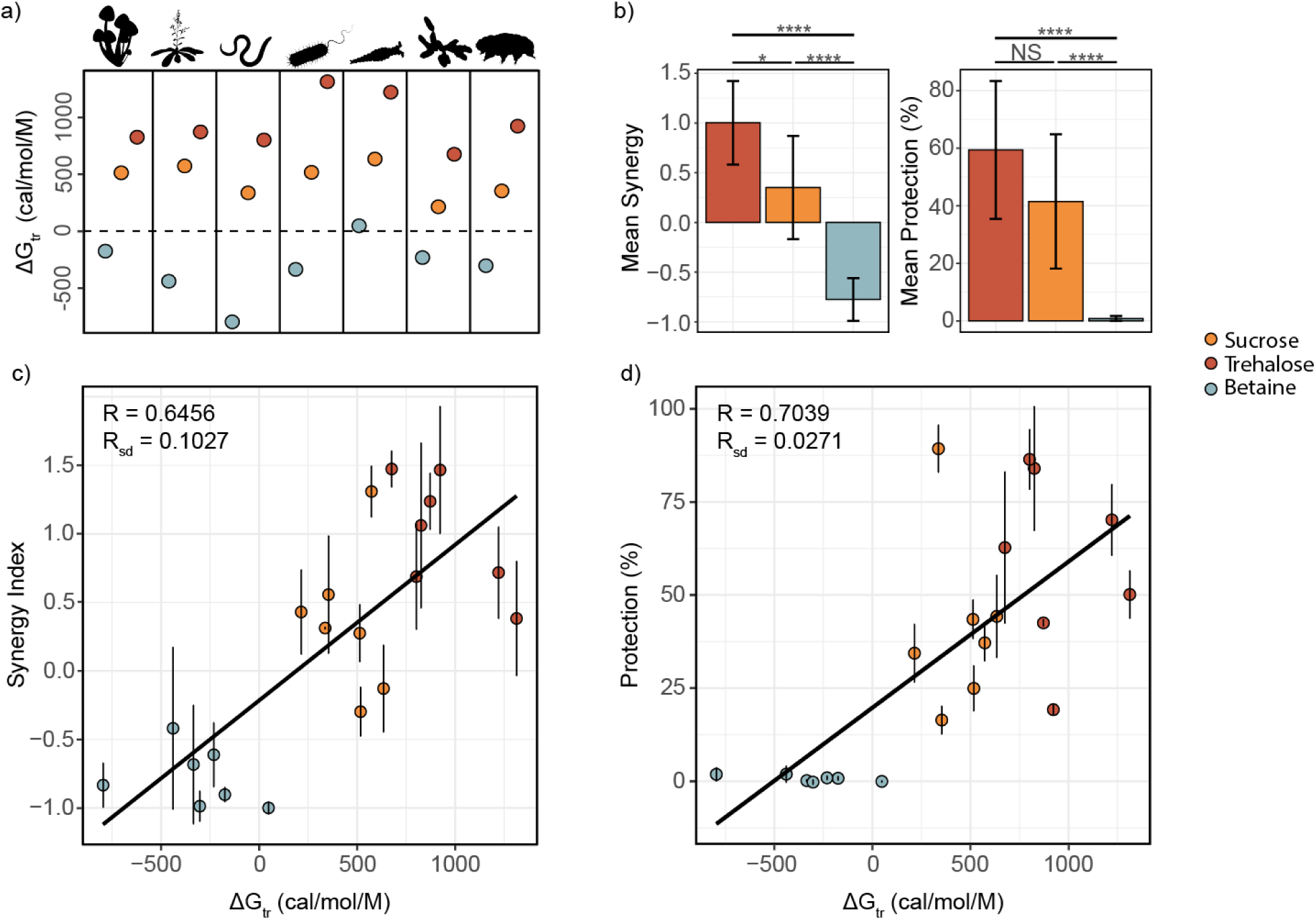
Transfer free energy acts as a predictor of synergy. **a)** The *ΔG_tr_* between each LEA_4 motif and each of the three selected cosolutes. **b)** The mean synergy (left) and mean protection (right) of LEA_4 motifs mixed with each of the three selected cosolutes. p-values determined using a two-way student’s t-test in R 4.3.0. **c)** A scatterplot correlating *ΔG_tr_* and synergy. **c)** A scatterplot correlating *ΔG_tr_* and percent protection. For all correlations, R (correlation coefficient) was generated using a Pearson’s Correlation in R 4.3.0. R_sd_ (the standard deviation of the correlation coefficient) was generated using a Monte Carlo error simulation approach for correlations (see *Methods*).

The mean protection of mixtures of LEA_4 motifs and these three cosolutes follows the same trend as the TFE. Trehalose mixtures are on average more protective than sucrose mixtures, which are more protective than betaine mixtures (Fig. 5b). We see a similar trend present in the synergy of each cosolute. Trehalose and sucrose, which have a positive *ΔG_tr_* with LEA_4 motifs, tend to act synergistically (Fig. 5b). On the other hand betaine has a negative *ΔG_tr_* and only produces antagonistic effects in our CS synergy assay. (Fig. 5b). Correlating *ΔG_tr_* with synergy for each of these mixtures reveals a strong, statistically significant relationship (Fig. 5c). We see a similarly strong relationship between *ΔG_tr_* and protection (Fig. 5d). Taken together, these results imply a mechanism for cosolute induced synergy where synergistic cosolutes promote, and antagonistic cosolutes inhibit, the protein:protein interactions by LEA_4 motifs.

## Discussion

The role of IDPs and their motifs, as well as functional synergy, is an emerging paradigm in desiccation tolerance and the study of IDPs as a whole [10,24,25,28,33,63]. Here, we probed the protective capacity of LEA_4 motifs for citrate synthase (CS) during desiccation as well as functional synergy between LEA_4 motifs and desiccation-enriched cosolutes. We demonstrate that LEA_4 motifs protect CS on their own or in concert with synergistic cosolutes. We report that functional synergy correlates with the *ΔG_tr_* of the LEA_4 motif with the given cosolute, indicating that stabilizing protein:protein interactions is beneficial to the function of LEA_4 motifs. We speculate that this solution repulsiveness could affect LEA_4 motif behavior in a variety of ways, from driving homo-oligomerization to stabilizing motif:client interactions. Overall, this study contributes to the growing body of literature on IDP-cosolute interactions and improves our understanding of the role of LEA_4 motifs in desiccation tolerance.

### LEA_4 motif protection during desiccation is client-specific

In contrast to previous work that found LEA_4 motifs do not protect LDH during desiccation, our observation that LEA_4 motifs are sufficient to protect citrate synthase underscores that different mediators of desiccation tolerance may be optimized for the protection of specific clients and/or may work through distinct mechanisms [64]. A possible explanation for this differential protection is that not all enzymes respond to desiccation in the same way. While CS forms non-functional aggregates during desiccation [33], no such aggregation is observed for LDH which instead is thought to become nonfunctional due to misfolding or destabilization [65,66]. While full-length LEA_4 proteins may be able to mediate protection against desiccation induced protein destabilization and aggregation, their motifs may only be sufficient to prevent aggregation.

### The molecular shielding theory and functional synergy

Molecular shielding is a prominent theory invoked to explain the protective function of IDPs during desiccation, particularly in the context of IDP-mediated prevention of protein aggregation. This theory posits that IDPs may act as entropic springs or “shields’’ that essentially occupy space and prevent the association of aggregation-prone proteins during desiccation. As mentioned above, for LEA proteins, a structural shift from a highly disordered state to a more ordered one has been suggested, though never empirically shown, to promote molecular shielding [15]. Our results do not provide evidence that such a shift occurs under the conditions tested, at least with peptides of this length. These findings are consistent with our recent work on full-length LEA protein structure in the presence of cosolutes, which found that cosolutes do not strongly affect local or global structure in the hydrated state [25,28,67,68].

We had theorized that molecular shields that take up more space would be more effective at preventing protein dysfunction during desiccation. However, here we observed no correlation between the change in R_g_ of a LEA_4 motif and its protective synergy with cosolutes. Several factors may explain this result, namely that the experiments we performed only explored the structure of LEA_4 motifs in the aqueous state, and thus gave us no insight into what occurs during the intermediate stages of desiccation. Alternatively, it may simply be that global dimensions are one of several factors in determining the protectiveness of an IDP. For example, the degree to which a molecular shield can transiently interact with aggregation-prone clients likely also influences protective capacity.

In summary, we observe that cosolutes alter the global dimensions of LEA_4 motifs, but not their secondary structure. We further show that this change in the global dimensions is not sufficient, at least on its own, to explain synergy in our system.

### The role of ΔG_tr_ in Synergy

In this study, we used a computational approach to approximate *ΔG_tr_*, which measures the change in free energy of a LEA_4 motif’s structure that can be attributed to the presence of a cosolute. Using this technique, we observe a significant correlation between *ΔG_tr_* and synergy. This implies a general mechanism for synergy in which synergistic cosolutes stabilize protein:protein interactions in IDPs as opposed to stabilizing protein:solvent interactions.

Given the relative simplicity of our experimental system, we envision only three possible forms of protein:protein interactions that could be stabilized by repulsive cosolutes. First, one might see an increase in intra-protein interactions, leading to changes in a motif’s secondary structure, global dimensions, or both. Alternatively, one may see an increase in inter-protein interactions, either through the homotypic association of motifs with each other, or through heterotypic associations with CS.

Our CD spectroscopy and SAXS data indicate no significant change in the monomeric ensemble of LEA_4 motifs, other than global compaction which does not correlate with synergy. This would logically rule out intra-protein interactions being a driving force behind synergy. For this reason, we consider inter-protein interactions, rather than intra-protein, to be the most likely mechanistic drivers of synergy. Furthermore, the lack of an increase in a motif’s R_g_ in the presence of cosolutes makes it unlikely that homotypic oligomerization is taking place on a meaningful scale. This leaves us with a single mechanism, the stabilization of heterotypic oligomerization between motifs and CS, that we feel is the most parsimonious to explain synergy between LEA_4 motifs and cosolutes. This idea directly conflicts with the idea that LEA_4 proteins and their motifs are simply inert crowders, and implies the necessity of direct interactions between LEA_4 motifs and their desiccation-sensitive clients. Future empirical studies quantifying changes in motif:CS interactions could be aimed at verifying this assertion.

## Conclusion

Beyond desiccation tolerance, intrinsically disordered proteins are ubiquitous across life and make up a significant fraction of the proteome of most organisms [36]. A large portion of IDPs contain short but important functional elements known as Short Linear Motifs (SLiMs) [69]. Here, we demonstrate that motifs from a family of desiccation-related IDPs, LEA_4 proteins, sense their chemical environment, causing them to undergo structural and functional changes in the process. Furthermore, we show that transfer free energy can be used to explain the influence observed between these motifs and their chemical environment. This suggests a model for functional synergy in which cosolutes lower the energetic cost of a protein adopting an optimal conformation. This research informs our understanding of the importance of LEA_4 motifs in desiccation tolerance. It also contributes to the growing body of literature documenting functional synergy between IDPs and cosolutes during desiccation [23–25,28,63]. More broadly, our work provides evidence that TFEs for different cosolutes can be used to approximate its ability to exact changes in an IDP or IDR’s ensemble and function.

## Methods

### LEA_4 Motifs Consensus Sequence and Cosolute Identification

The LEA_4 motif sequence was obtained from pfam and used as a query in a BLAST analysis against proteomes from different organisms [70–72]. Matching sequences (see File S1 for accession numbers) were selected, and LEA_4 motifs were identified via Pfam searches [70]. LLEA_4 motif sequences for each protein were aligned using ClustalX, and alignments were visualized using WebLogo (https://weblogo.berkeley.edu/logo.cgi) [73]. Selection of cosolutes was based on previous reports in the literature (Table 1).

### Citrate Synthase Assay

The Citrate Synthase Kit (Sigma-Aldrich Cat. CS0720) was adapted for use in this assay. All samples were prepared in triplicate. Lyophilized peptides were resuspended in either purified water (for samples to be desiccated) or 1X assay buffer (for control samples) to a concentration of ~20 g/L, and diluted as necessary for lower concentrations. Porcine citrate synthase (Sigma-Aldrich Cat. C3260, UniProt P00889) was added at a ratio of 1:10 to the resuspended protectants. Non-desiccated control samples were measured according to the assay kit instructions immediately after resuspension. Desiccated samples were subjected to six rounds of desiccation and rehydration (1 hour speedvac desiccation [SAVANT, SpeedVac SC110] followed by resuspension in water). After the final round of desiccation, samples were resuspended in 10 μL of cold 1X assay buffer. Samples were diluted 1:100 in the assay reaction mixture supplied, and all subsequent steps followed the kit instructions. The colorimetric reaction was measured for 90 seconds at 412 nm using the Spark 10M (Tecan).

### Citrate Synthase Assay - Assessing Functional Synergy

Synergy experiments were performed using an adapted version of the above protocol. To assess the synergy of a simple motif-cosolute combination, three solutions were prepared and assayed (in triplicate): one containing 0.1 mg/mL of a LEA_4 motif, one containing 1.5 mM of a cosolute, and one containing a mixture of the two at the same concentrations. These samples were desiccated side-by-side under identical conditions before being reconstituted and assayed using the same reagents. This generates three protective capacity values which are sufficient to calculate the “synergy index” (below).

### Calculation of Synergy Index

In the assay described above, the efficiency of protectants was assessed independently and as a mixture. A prediction for the protective capacity of a peptide/cosolute mixture was calculated by simply taking the sum of their individual protections.

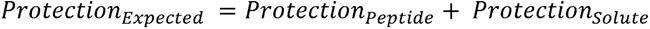

For statistical purposes, we calculated the standard error (SE) of this value using the equation below.

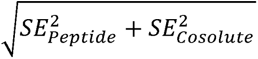

The synergy index was then calculated using the formula for percent error.

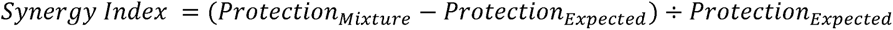

### Small Angle X-Ray Scattering (SAXS)

All SAXS measurements were performed by the SIBYLS group at the Lawrence Berkeley National Laboratory HT-SAXS beamline [49–52]. Lyophilized peptides were dissolved in a buffer containing 100 mM NaCl and 20 mM sodium acetate (pH = 7.4) at concentrations of 4, 6, and 8 g/L. For mixed samples, cosolutes were dissolved at 0.438 M or until fully saturated before the addition of peptide. In accordance with SIBYLS guidelines, blank buffer samples were provided for each protein/cosolute mixture. All data reported are on peptides at 6 g/L, as this concentration generally yielded a strong signal with little noise.

### Guinier Analysis

Buffer subtractions and Guinier analysis were performed using BioXTAS RAW v. 2.1.4 [74,75]. Because of the intrinsically disordered nature of the LEA_4 motifs, a q_max_R_g_ of 1.1 was used to establish linear fits in the Guinier region. Guinier regions showing characteristic evidence of sample degradation, solution repulsion, or aggregation were excluded from all analyses and plots.

### Kratky Analysis

Kratky plots were generated from buffer-subtracted .dat files in R Studio version 2023.03.0+386 using R v.4.3.1. First, the .dat file was processed into a .csv file containing only the raw q and I(q) values. Then, the values were displayed as a scatterplot where the x-axis was q and the y-axis was I(q) * q^2^. A csv copy of each scattering profile is available in supplementary data (File S1).

### CD Spectroscopy

Lyophilized LEA_4 motifs were resuspended in a buffer containing 25 mM tris (pH = 7) to a concentration of 100 μM. For samples measured with a cosolute, the lyophilized peptides were resuspended in a solution containing Tris buffer and 100 mM of the cosolute.. Peptide concentrations were quantified using Qubit™ Protein Assay (Catalog number Q33211, Thermo Fisher Scientific, USA). Aliquots of the peptide suspensions were measured in a 1 mm quartz cuvette in a circular dichroism spectrometer (Jasco, J-1500 model, Japan). Raw spectra are provided in supplementary data (File S1).

### CD Deconvolution

CD spectra were deconvolved using Beta Structure Selection method (BeStSel) [53,54]. The raw data for each CD spectrum, measured in molar residue ellipticity, were pasted into the online portal and analyzed using the ‘Single spectrum analysis’ setting.. Only data from 190 nm to 250 nm was considered in the analysis.

CD spectra were then deconvolved using DichroWeb [55,56]. Each spectra was uploaded to DichroWeb and analyzed using the SELCON3 algorithm, which considers CD signals between 190 nm and 240 nm. Secondary structure annotation was determined via comparison with the “SET7” dataset, which contains both folded and denatured proteins [56].

### Transfer Free Energy (TFE) Calculations

To calculate *ΔG_tr_*, we used empirical TFE values from Auton and Bolen 2005 and Hong 2015 [57,62]. Using these values, *ΔG_tr_* can be calculated using the following equation:

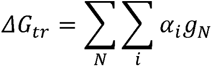

Here, *ΔG_tr_* is the change in free energy undergone by a protein of fixed conformation upon transfer from water to a 1 M cosolute solution, N is the chemical group, i is a numerical index for all instances of the chemical group, *α* is the relative solvent accessibility of the chemical group as a fraction of 1, and g is the experimental value of the transfer free energy for that chemical group. Because of the inherently disordered nature LEA_4 motifs, we assume *α* = 1 for all protein residues.

### Monte Carlo Correlation Error Simulations

To generate the correlation statistics in figures 3 and 4, custom code was written to test a large variety of correlations with simulated error. Two normal distributions were created for each datapoint, one using the x-axis mean and standard deviation, and the other using the y-axis mean and standard deviation. By generating random values from the resulting normal distributions, we could simulate the effect of random error on each point in both the x and y direction*. Using this system on each of our correlation plots, we performed 100,000 error simulations, each having a unique Pearson correlation coefficient, R. We can use this list of R values to calculate R_sd_. The code used to execute this method is available in supplementary material (File S2) and is well annotated.

*Our data pertaining to CD deconvolution had no error; thus, the standard deviation was set to 0.

## Supplementary Material Descriptions

File S1: Contains raw .xlsx and .csv files that store the data used in this work, organized by figure. Also contains SAXS scattering profiles and CD spectra, where applicable.

File S2: Contains all custom code, including R scripts for the generation of figures and Python scripts for performing TFE computations.

## Abbreviations

IDPs: Intrinsically Disordered Proteins
LEA: Late Embryogenesis Abundant
CS: Citrate Synthase
LDH: Lactate Dehydrogenase
SAXS: Small Angle X-Ray Scattering
R_g_: Radius of Gyration
CD: Circular Dichroism
TFE: Transfer Free Energy

## Acknowledgements

Support for this project came from the NSF via the IntBio research program under awards 2128069 to TCB, 2128067 to SS, and 2128068 to ASH. Graduate students from the University of Wyoming were supported in part by the USDA National Institute of Food and Agriculture, Hatch project #1012152. We thank members of the Water and Life Interface Institute (WALII), supported by NSF DBI grant #2213983, for helpful discussions.

We thank Dr. Greg Hura and Kathryn Burnett for their correspondence and help in performing the SAXS experiments. SAXS experiments were conducted at the Advanced Light Source (ALS), operated by Lawrence Berkeley National Laboratory on behalf of the Department of Energy, Office of Basic Energy Sciences, through the Integrated Diffraction Analysis Technologies (IDAT) program, supported by DOE Office of Biological and Environmental Research.

## Author Contributions

Conceptualization: **VN, KN, ASH, SS, TCB**

Data Curation: **VN, KN, EG, MM, FY**

Formal Analysis: **VN**

Funding Acquisition: **ASH, SS, TCB**

Investigation: **VN, EG, FY**

Citrate synthase assay: **VN, KN**

Circular dichroism spectroscopy: **EG, MM, FY**

Small angle X-ray scattering: **VN, FY**

TFE computation/analysis: **VN**

Methodology: **VN, KN, EG, FY**

Supervision: **ASH, SS, TCB**

Visualization: **VN, ASH, TCB**

Writing – Original Draft: **VN, TCB**

Writing – Review & Editing: **VN, KN, EG, MM, FY, ASH, SS, TCB**

---

**Figure S1.**
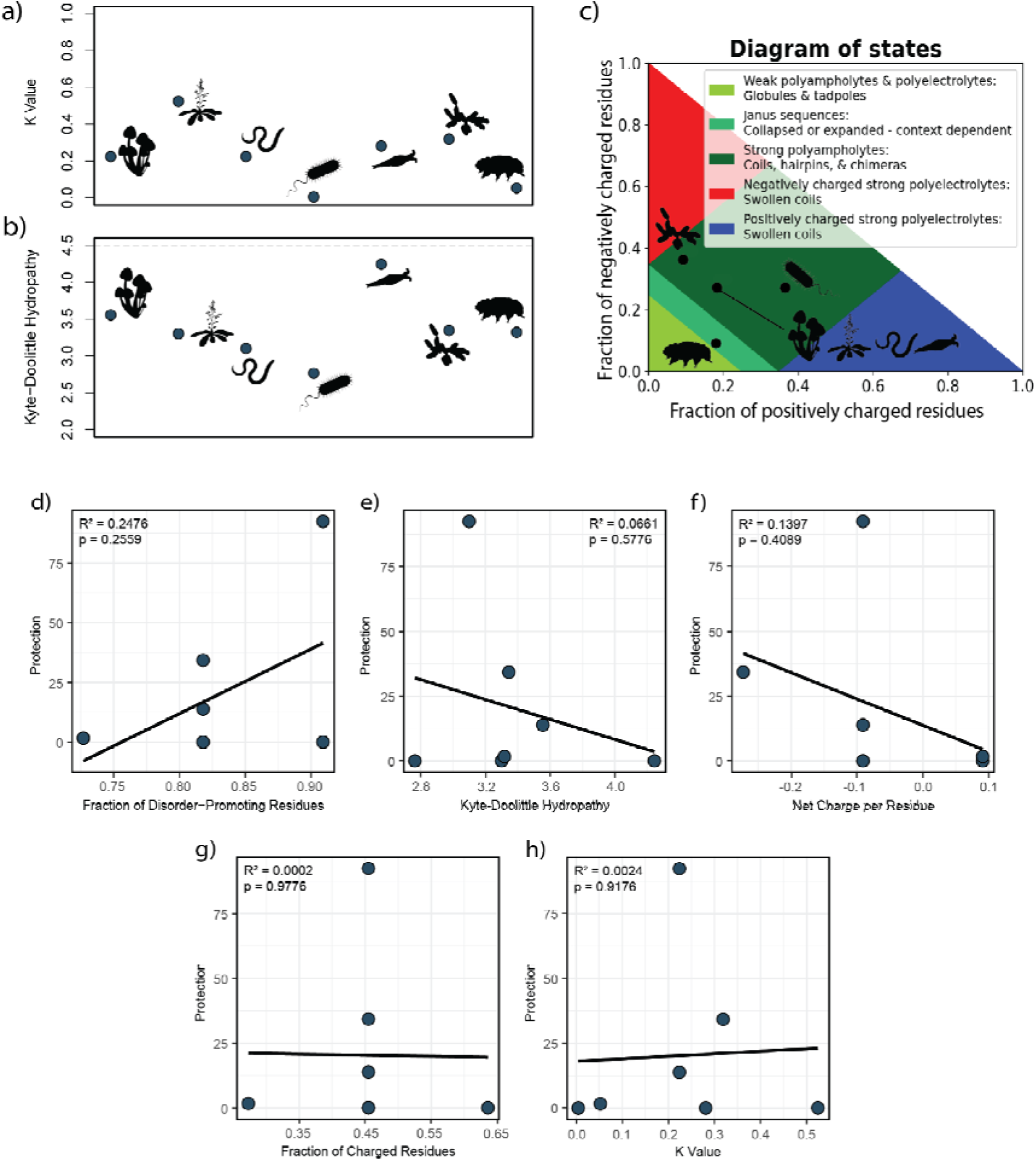
LEA_4 motifs from different organisms have diverse sequence features. **a)** The κ value (an approximation of the charge patterning) for each LEA_4 motif. Low values represent even charge patterning, and high values represent charge segregation. **b)** The Kyte-Doolittle hydropathy of each LEA_4 motif on a 0-9 scale. Higher values indicate more hydrophobicity. Dotted line represents “neutral” hydropathy, in which the peptide is neither hydrophilic nor hydrophobic. **c)** A Das-Pappu phase diagram of LEA_4 motifs, indicating their highly charged nature and sequence diversity. **d-h)** Scatter plots of a peptide’s sequence parameters vs its protection at 1.68 mM. **d)** Fraction of disorder-promoting residues. **e)** Kyte-Doolittle Hydropathy. **f)** Net Charge per Residue. **g)** Fraction of Charged Residues. **h)** Kappa value. R^2^ and p-values were generated using a Pearson’s Correlation in R 4.3.0.

**Figure S2.**
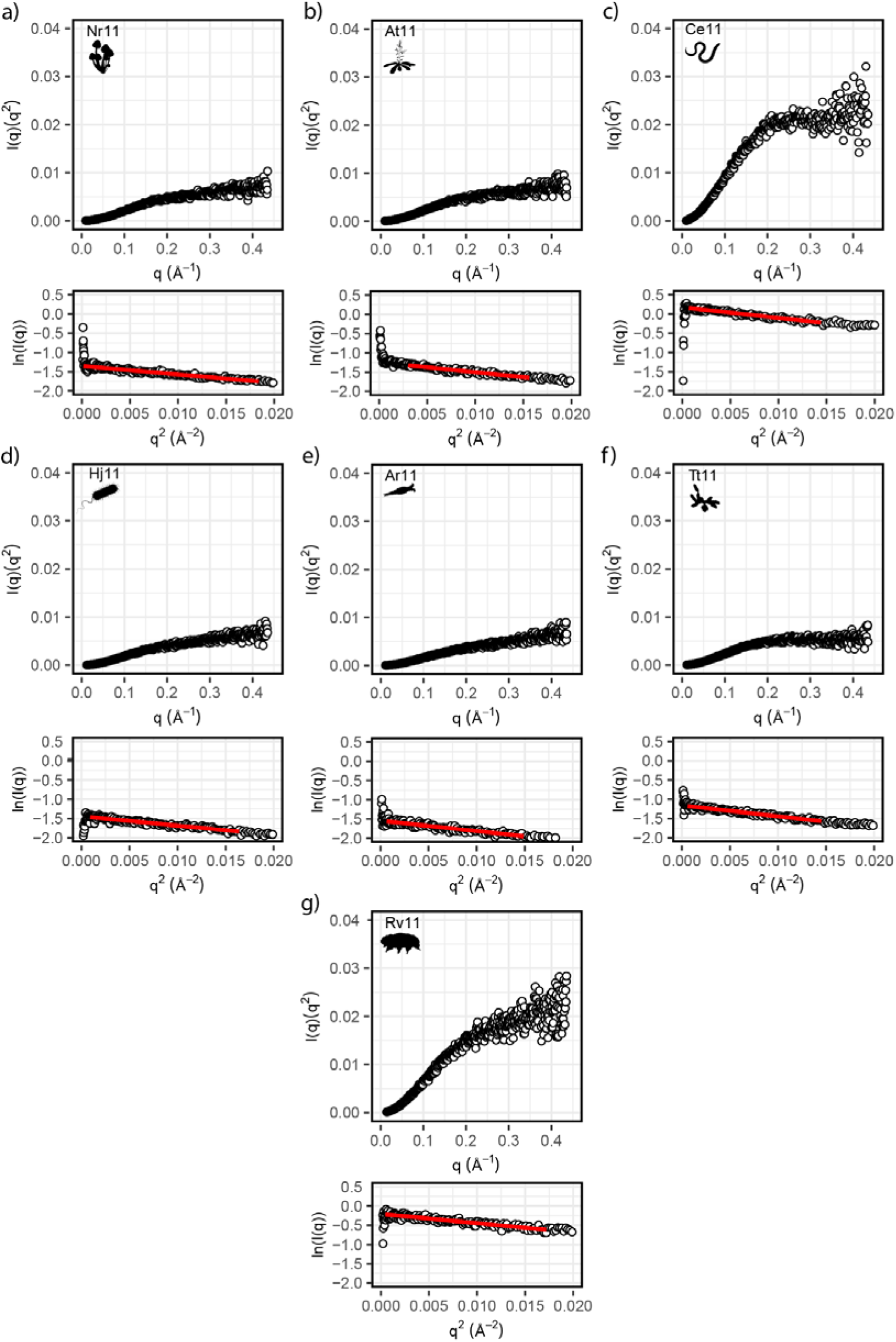
Representative HT-SAXS data of each LEA_4 motif in buffer, displayed as a Kratky plot (top) and a Guinier plot with a red linear fit (bottom). **a)** Nr11, **b)** At11, **c)** Ce11, **d)** Hj11, **e)** Ar11, **f)** Tt11, **g)** Rv11.

**Figure S3.**
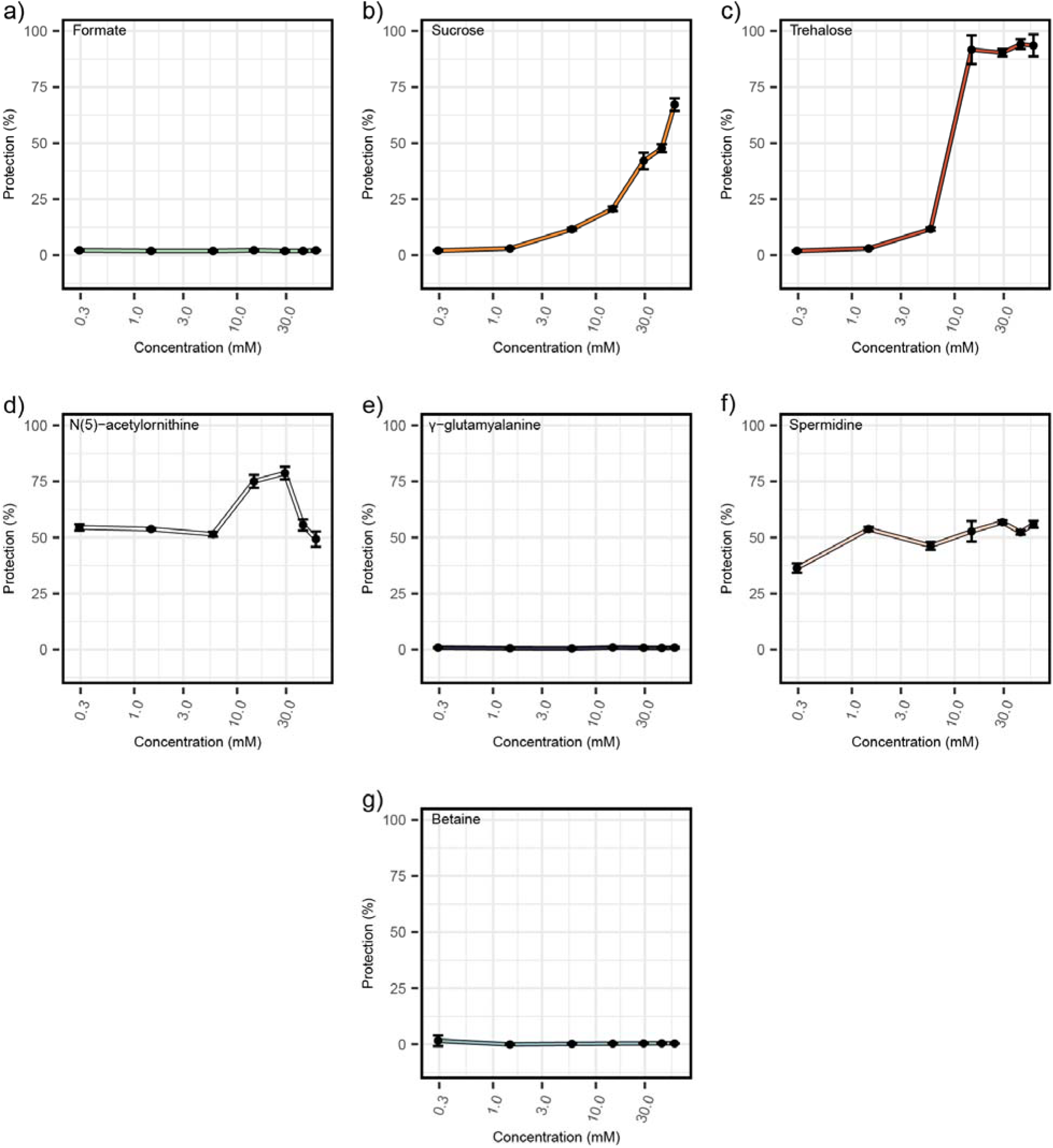
The ability of drying-enriched cosolutes to protect citrate synthase from aggregating during desiccation. **a)** Formate. **b)** Sucrose. **c)** Trehalose. **d)** N(5)-acetylornithine. **e)** γ-Glutamylalanine. **f)** Spermidine. **g)** Betaine.

**Figure S4.**
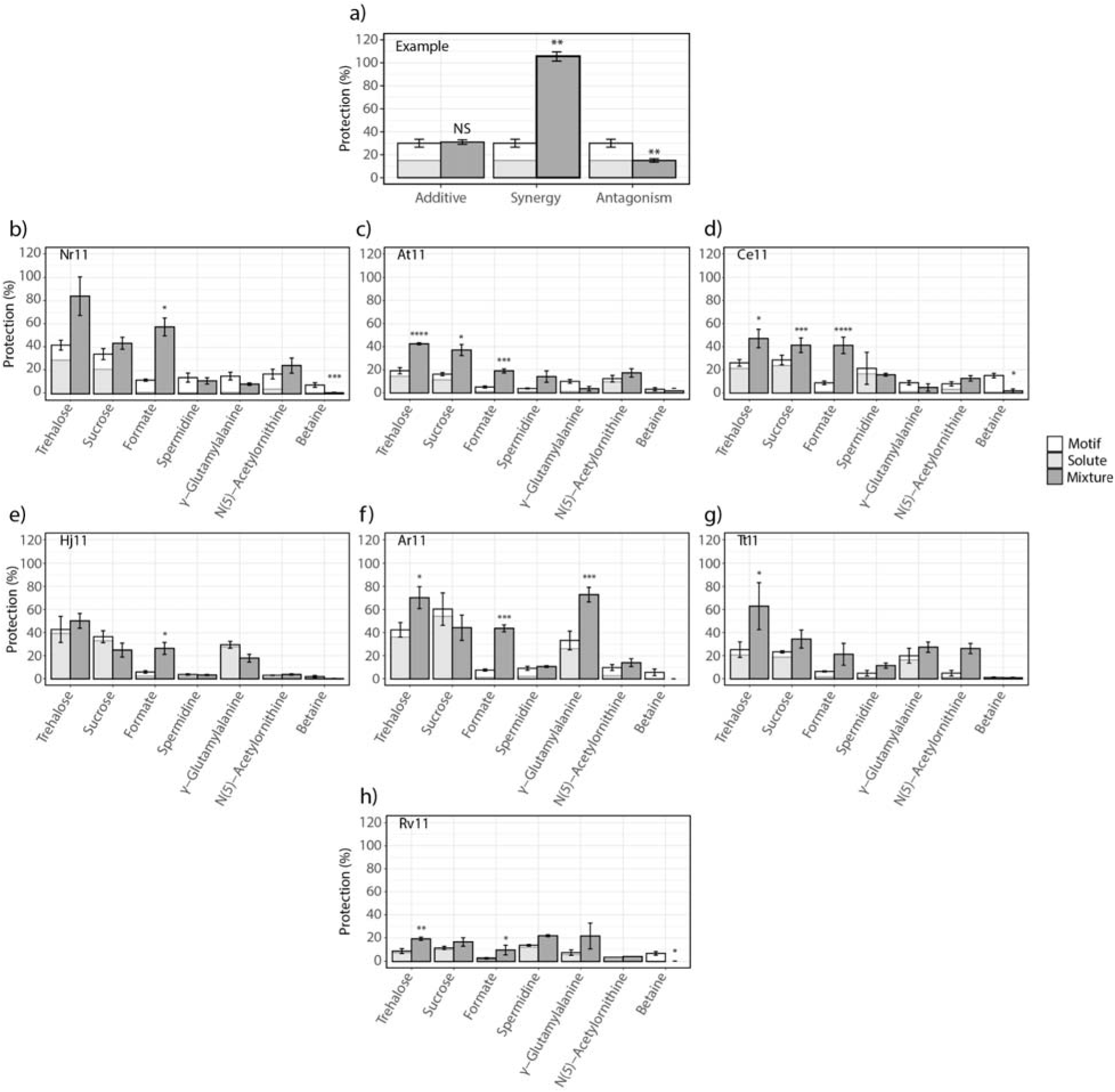
The ability of various LEA_4 motifs to protect in the citrate synthase assay. **a)** An example plot showing a cosolute that has no effect on protection (left), a cosolute that is synergistic (middle), and a cosolute that is antagonistic (right). **b)** Nr11, **c)** At11, **d)** Ce11, **e)** Hj11, **f)** Ar11, **g)** Tt11, **h)** Rv11. Error bars represent the standard error of each sample. All data was collected in triplicate. All statistical analysis is performed using a one-way Student’s t-test.

**Figure S5.**
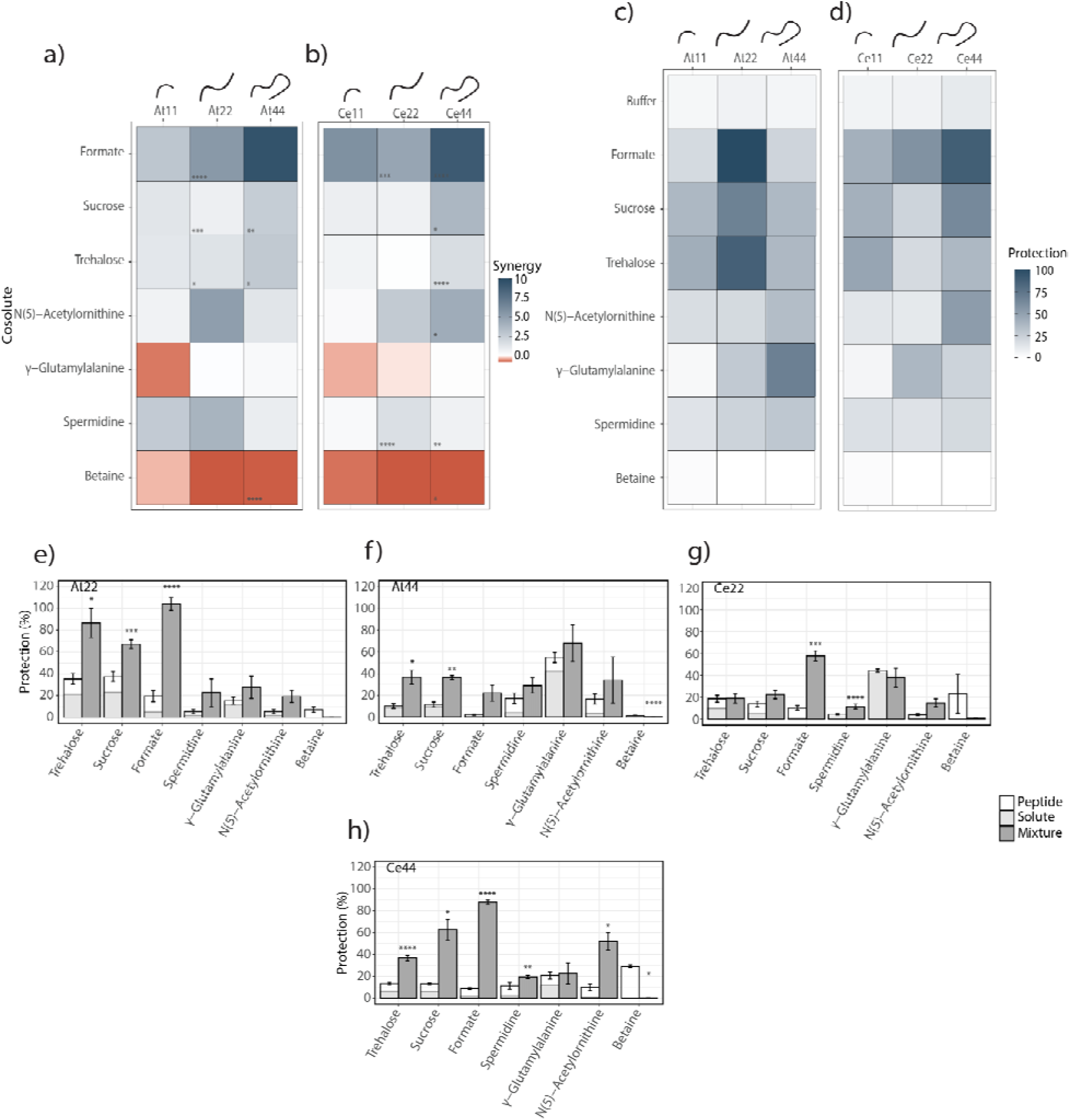
**a)** Heatmap of synergy index between *Arabidopsis thaliana* LEA_4 motifs of variable length and various cosolutes. **b)** Heatmap of synergy index between *C. elegans* LEA_4 motifs of variable length and various cosolutes. **c)** Heatmap of the protective capacity of mixtures of *Arabidopsis thaliana* LEA_4 motifs and various cosolutes. **d)** Heatmap of the protective capacity of mixtures of *C. elegans* LEA_4 motifs and various cosolutes. **e-h)** The ability of 2x and 4x LEA_4 motifs to protect in the citrate synthase assay. **e)** At22. **f)** At44. **g)** Ce22. **h**) Ce44. Error bars represent the standard error of each sample. All data was collected in triplicate. All statistical analysis is performed using a one-way Student’s t-test.

**Figure S6.**
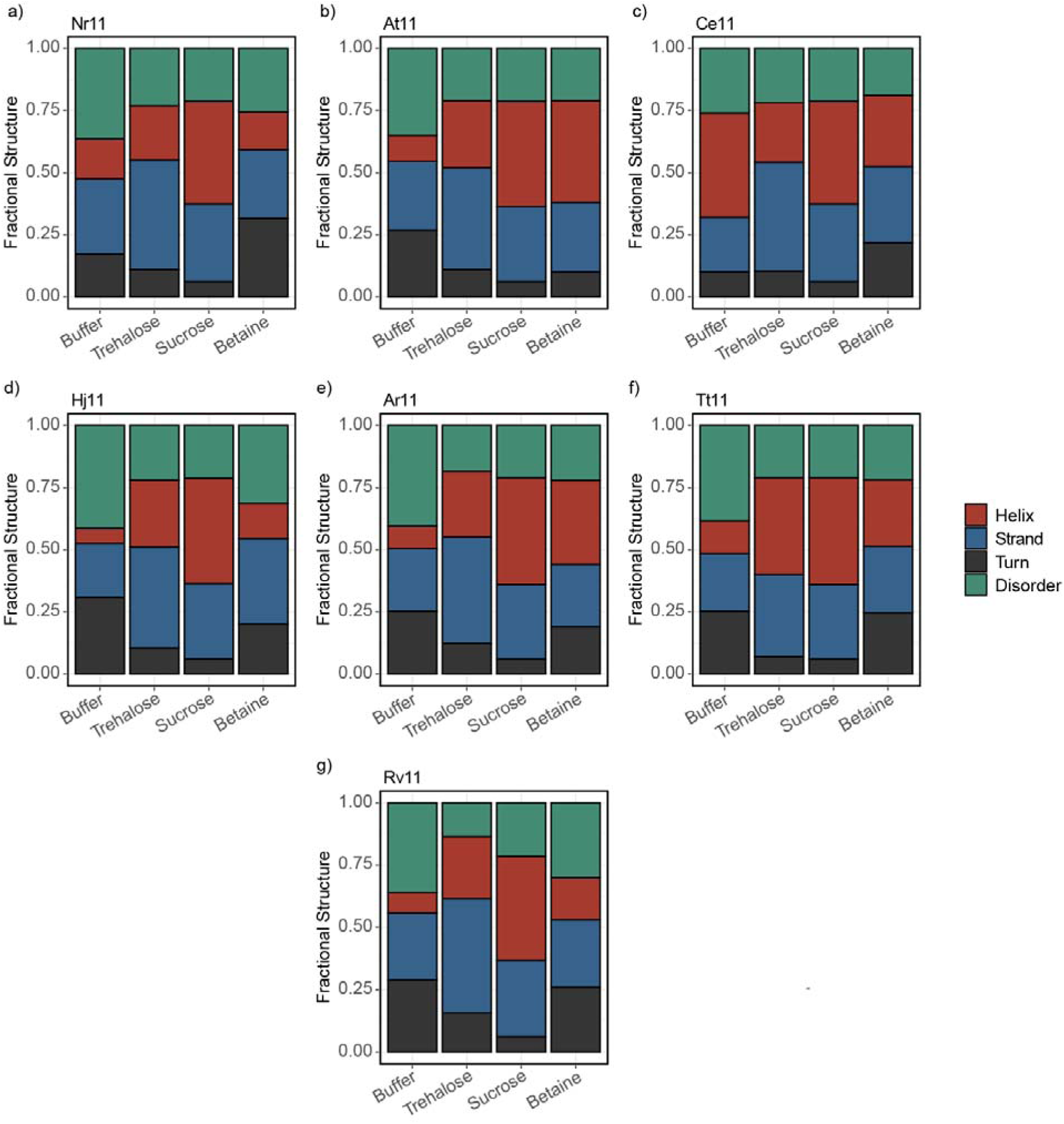
Deconvoluted fractional secondary structure from DichroWeb. **a)** Nr11, **b)** At11, **c)** Ce11, **d)** Hj11, **e)** Ar11, **f)** Tt11, **g)** Rv11.

