## Supplementary material for "LEA_4 motifs function alone and in conjunction with synergistic cosolutes to protect a labile enzyme during desiccation": File S2: PD50.R

```
library(readxl)
library(drc)
library(tidyverse)
library(patchwork)
library(ggborderline)

setwd("C:/Users/Vincent/OneDrive/Documents/Boothby Lab Work/Motif Synergy Project/CS Assay")

type <- "Peptide"

file <- read_excel(paste(type, "PD50.xlsx"))

# x0 <- rev(temp$Concentration)
# y0 <- rev(temp$Protections)
# e0 <- rev(temp$Stdevs)

# fit <- drm(y0~x0, data = data.frame(x0, y0), fct = LL.4())

# PD50 = 58.4
# acc = 0
# 
# for(i in seq(0, 100, length.out = 10000))
# {
#   p = as.numeric(predict(fit, data.frame(i, i)))
#   if((abs(50 - acc) >= abs(50 - p)))
#   {
#     PD50 = i
#     acc = p
#   }
# }
# 
# PD50

if(type == "Metabolite")
{
  protectants <- c("Formate", "Sucrose", "Trehalose", "Na-acetylornithine", "y-glutamyalanine", "Spermidine", "Betaine")
  col_list <- c("#A3C6A8", "#F28F3B", "#C8553D", "#FFFFFF", "#2D3047", "#FFD5C2", "#93B7BE")
  barsize <- 0.1
}
if(type == "Peptide")
{
  protectants <- c("Anaerobic fungus", "Arabadopsis", "C. elegans", "Halobecteria", "Rotifer", "Thermophilic archae", "Tardigrade")
  col_list <- c("#284b63", "#284b63", "#284b63", "#284b63", "#284b63", "#284b63", "#284b63")
  barsize <- 0.1
}

plot_list <- c()

for(i in 1:length(protectants))
{
  if(type == "Metabolite")
  {
    temp <- subset(file, Cosolutes == protectants[i])
  }
  if(type == "Peptide")
  {
    temp <- subset(file, Species == protectants[i])
  }
  
  temp$Concentration <- round(temp$Concentration, digits = 3)
  
  plot_list[[i]] <- ggplot(data = temp) +
    geom_borderline(stat = "identity", aes(x = Concentration, y = Protections), size = 0.5, color = col_list[i], lwd = 1, bordercolor = "black") +
    geom_point(aes(x = Concentration, y = Protections), size = 1) +
    geom_errorbar(aes(x = Concentration, ymin = Protections - Stdevs, ymax = Protections + Stdevs, width = barsize), color = "black") +
    theme_bw()+
    theme(
      plot.title = element_text(hjust = 0.05, size = 6, vjust = -10),
      axis.title = element_text(size =6),
      axis.text.y = element_text(size =6),
      axis.text.x = element_text(size = 6, angle = 70, vjust = 1, hjust = 1),
      legend.text=element_text(size=6),
      legend.position = 'none', aspect.ratio = 1,
      legend.title = element_blank(),
      panel.border = element_rect(colour = "black", fill=NA, size=1)) +
    labs(x = "Concentration (mM)", y = "Protection (%)", title = protectants[i]) +
    scale_x_log10() +
    ylim(-10, 100)
}

(plot_list[[1]] | plot_list [[2]] | plot_list[[3]]) / (plot_list[[4]] | plot_list [[5]] | plot_list[[6]]) / (plot_list[[7]] | plot_list [[7]] | plot_list[[7]])
```
