## Supplementary material for "LEA_4 motifs function alone and in conjunction with synergistic cosolutes to protect a labile enzyme during desiccation": File S2: Heatmap.R

```
library(readr)
library(dplyr)
library(ggplot2)
library(readxl)
library(ggpmisc)

setwd("C:/Users/Vincent/OneDrive/Documents/Boothby Lab Work/Motif Synergy Project/CS Assay")

file <- read_excel('Heatmap Master.xlsx', sheet = '11mers')

file$Solute = factor(file$Solute, levels = c("Betaine", "Spermidine", "y-Glutamylalanine", "No-Acetylornithine", "Trehalose", "Sucrose", "Formate", "Buffer"))

#Include for synergy heatmaps, comment out for protection ones
file <- subset(file, Solute != "Buffer")

heat <- ggplot(data = file, aes(x = Peptide, y = Solute, fill = Synergy)) +
  scale_fill_gradient2(
    low = "#C8553D",
    mid = "white",
    high = "#284b64",
    aesthetics = "fill",
    guide = "colorbar",
    limits = c(0, 6)) +
  geom_tile(color = "black", show.legend = T) +
  scale_x_discrete(position = "top") +
  coord_equal() +
  theme_bw()

heat

heat <- ggplot(data = file, aes(x = Peptide, y = Solute, fill = Synergy)) +
  scale_fill_gradient2(
    low = "#C8553D",
    mid = "white",
    high = "#284b64",
    aesthetics = "fill",
    guide = "colorbar",
    limits = c(-1, 0)) +
  geom_tile(color = "black", show.legend = T) +
  scale_x_discrete(position = "top") +
  coord_equal() +
  theme_bw()

heat
```
