## Supplementary material for "LEA_4 motifs function alone and in conjunction with synergistic cosolutes to protect a labile enzyme during desiccation": File S2: Cor Script.R

```
library(tidyverse)
library(readxl)
library(ggpmisc)

setwd("C:/Users/Vincent/OneDrive/Documents/Boothby Lab Work/Motif Synergy Project")

file <- read_excel('Data Master.xlsx')

# ttmers <- subset(file, Peptide == "At-22mer" | Peptide == "Ce-22mer")
# ffmers <- subset(file, Peptide == "At-44mer" | Peptide == "Ce-44mer")
file <- subset(file, Peptide == "Af-11mer" | Peptide == "At-11mer" | Peptide == "Ce-11mer" | Peptide == "Hb-11mer" | Peptide == "Rt-11mer" | Peptide == "Ta-11mer" | Peptide == "Tg-11mer")

file <- subset(file, Solute != "Buffer")
temp <- file
x = file$`ΔG`
y = file$`Protection`

if (x[1] == file$'ΔG'[1])
{
  temp <- subset(file, Solute == "Sucrose" | Solute == "Trehalose" | Solute == "Betaine")
  xlims = c(-200,200)
  xlabs = "ΔG (cal/mol/M)"
  x = temp$'ΔG'
  cols <- c("#93B7BE", "#F28F3B", "#C8553D")
}
if (x[1] == file$'Rg'[1])
{
  temp <- subset(file, Solute != "Buffer")
  temp <- subset(temp, SAXS_Clean == "Y")
  xlims = c(0, 10)
  xlabs = "Rg"
  x = temp$'Rg'
  xerr = temp$'Rg_StDev'
  cols <- c("#93B7BE", "#6A8470", "#FFFFFF", "#FFD5C2", "#F28F3B", "#C8553D", "#2D3047")
}
if (x[1] == file$'Rg_Delta'[1])
{
  temp <- subset(file, Solute != "Buffer")
  temp <- subset(temp, SAXS_Clean == "Y")
  xlims = c(10,-10)
  xlabs = "Change in Rg (Å)"
  x = temp$'Rg_Delta'
  xerr = temp$'Rg_StDev'
  cols <- c("#93B7BE", "#6A8470", "#FFFFFF", "#FFD5C2", "#F28F3B", "#C8553D", "#2D3047")
}
if (x[1] == file$'Rg_Percent'[1])
{
  temp <- subset(file, Solute != "Buffer")
  temp <- subset(temp, SAXS_Clean == "Y")
  xlims = c(0, -0.4)
  xlabs = "Fractional dRg"
  x = temp$'Rg_Percent'
  cols <- c("#93B7BE", "#6A8470", "#FFFFFF", "#FFD5C2", "#F28F3B", "#C8553D", "#2D3047")
}
if (y[1] == file$`Synergy`[1])
{
  ylims = c(-1, 2)
  ylabs = "Synergy Index"
  y = temp$'Synergy'
  yerr = temp$'Syn_Error'
}
if (y[1] == file$Protection[1])
{
  ylims = c(0, 110)
  ylabs = "Protection (%)"
  y = temp$'Protection'
  yerr = temp$'Prot_Error'
}

data <- data.frame(x,y)
data <- cbind(data, temp$Solute)

ggplot(data = data) +
  geom_point(aes(x = x, y = y, fill = temp$Solute), size = 2.5, color = "black", shape = 21) +
  scale_fill_manual(values = cols) +
  geom_errorbar(data = data, aes(x = x, ymin = y-yerr, ymax = y+yerr), lwd = 0.4) +
  # geom_errorbar(data = data, aes(y = y, xmin = x-xerr, xmax = x+xerr), lwd = 0.4) +
  geom_smooth(method = lm, aes(x = x, y = y), se = F, color = "black") +
  theme_bw()+
  theme(
    plot.title = element_text(hjust = 0.05, size = 14, vjust = -8),
    axis.title = element_text(size =14),
    axis.text.y = element_text(size =10),
    axis.text.x = element_text(size = 10),
    legend.text=element_text(size=14),
    legend.position = 'none', aspect.ratio = 1,
    legend.title = element_blank(),
    panel.border = element_rect(colour = "black", fill=NA, size=1)) +
  labs(x = xlabs, y = ylabs)
  
cor <- cor.test(x, y, method = "pearson")
r <- unname(cor$estimate)
r^2
cor$p.value
```
