## Supplementary material for "LEA_4 motifs function alone and in conjunction with synergistic cosolutes to protect a labile enzyme during desiccation": File S2: Short Stripchart.R

```
library(readr)
library(dplyr)
library(ggplot2)
library(readxl)
library(ggpmisc)

setwd("C:/Users/Vincent/OneDrive/Documents/Boothby Lab Work/Motif Synergy Project/SAXS")

file <- read_excel('Clean SAXS Data.xlsx')

file <- subset(file, Peptide != "At-22mer" & Peptide != "At-44mer" & Peptide != "Ce-22mer" & Peptide != "Ce-44mer")
file$Peptide <- factor(file$Peptide, levels = c("Af-11mer", "At-11mer", "Ce-11mer", "Hb-11mer", "Rt-11mer", "Ta-11mer", "Tg-11mer"), ordered = T)
file <- subset(file, Concentration == 6)
short <- subset(file, Solute != "Buffer")

buff <- subset(file, Solute == "Buffer")
norm_list <- buff$Rg

Solutes <- unique(short$Solute)
Peptides <- c("Af-11mer", "At-11mer", "Ce-11mer", "Hb-11mer", "Rt-11mer", "Ta-11mer", "Tg-11mer")

Lines <- c(1.5, 2.5, 3.5, 4.5, 5.5, 6.5)

cols <- c()
Solute <- file$Solute

for(i in 1:length(file$Solute))
{
  if(Solute[i] == "Formate")
  {
    cols <- append(cols, "#588B8B")
  }else if (Solute[i] == "No-Acetylornithine")
  {
    cols <- append(cols, "#FFFFFF")
  }else if (Solute[i] == "Spermidine")
  {
    cols <- append(cols, "#FFD5C2")
  }else if (Solute[i] == "Sucrose")
  {
    cols <- append(cols, "#F28F3B")
  }else if (Solute[i] == "Trehalose")
  {
    cols <- append(cols, "#C8553D")
  }else if (Solute[i] == "y-Glutamylalanine")
  {
    cols <- append(cols, "#2D3047")
  }
}

ggplot(data = short) +
  geom_errorbar(position = position_dodge(0.75), aes(x = Peptide, ymin = Rg-Rg_StDev, ymax = Rg+Rg_StDev, fill = Solute), color = "black", width = 0, lwd = 1) +
  geom_point(position = position_dodge(0.75), aes(x = Peptide, y = Rg, fill = Solute), size = 3, pch = 21, color = "black") +
  geom_segment(aes(x = 0, xend = 1.5, y = norm_list[1], yend = norm_list[1]), lty = "dashed", size = 0.5) +
  geom_segment(aes(x = 1.5, xend = 2.5, y = norm_list[2], yend = norm_list[2]), lty = "dashed", size = 0.5) +
  geom_segment(aes(x = 2.5, xend = 3.5, y = norm_list[3], yend = norm_list[3]), lty = "dashed", size = 0.5) +
  geom_segment(aes(x = 3.5, xend = 4.5, y = norm_list[4], yend = norm_list[4]), lty = "dashed", size = 0.5) +
  geom_segment(aes(x = 4.5, xend = 5.5, y = norm_list[5], yend = norm_list[5]), lty = "dashed", size = 0.5) +
  geom_segment(aes(x = 5.5, xend = 6.5, y = norm_list[6], yend = norm_list[6]), lty = "dashed", size = 0.5) +
  geom_segment(aes(x = 6.5, xend = 8, y = norm_list[7], yend = norm_list[7]), lty = "dashed", size = 0.5) +
  geom_vline(xintercept=c(1.5, 2.5, 3.5, 4.5, 5.5, 6.5)) +
  scale_color_manual(values = c("#93B7BE", "#588B8B", "#FFFFFF", "#FFD5C2", "#F28F3B", "#C8553D", "#2D3047")) +
  scale_fill_manual(values = c("#93B7BE", "#588B8B", "#FFFFFF", "#FFD5C2", "#F28F3B", "#C8553D", "#2D3047")) +
  theme_bw() +
  theme(
    plot.title = element_text(hjust = 0.05, size = 14, vjust = -8),
    axis.title = element_text(size =14),
    axis.text.y = element_text(size =10),
    axis.text.x = element_text(size = 10),
    legend.text=element_text(size=14),
    legend.position = "none",
    aspect.ratio = 0.6,
    legend.title = element_blank(),
    panel.border = element_rect(colour = "black", fill=NA, size=1),
    panel.grid.major = element_blank(), 
    panel.grid.minor = element_blank()) +
  labs(y = "Rg (Å)")
```
