## Supplementary material for "LEA_4 motifs function alone and in conjunction with synergistic cosolutes to protect a labile enzyme during desiccation": File S2: CD Patchwork.R

```
library(tidyverse)
library(readxl)
library(ggpmisc)
library(patchwork)

setwd("C:/Users/Vincent/OneDrive/Documents/Boothby Lab Work/Motif Synergy Project/CD Spec")

file <- read_excel('CD Curves.xlsx', sheet = 'Ta-11mer')

Raw <- ggplot(data = file) +
  geom_line(aes(x = Wavelength, y = Buffer / 1000), size = 1.2, color = "grey") +
  geom_line(aes(x = Wavelength, y = Trehalose / 1000), size = 1.2, color = "#C8553D") +
  geom_line(aes(x = Wavelength, y = Sucrose / 1000), size = 1.2, color = "#F28F3B") +
  geom_line(aes(x = Wavelength, y = Betaine / 1000), size = 1.2, color = "#93B7BE") +
  theme_bw()+
  theme(
    plot.title = element_text(hjust = 0.05, size = 14, vjust = -8),
    axis.title = element_text(size =14),
    axis.text.y = element_text(size =10),
    axis.text.x = element_text(size = 10),
    legend.text=element_text(size=14),
    legend.position = 'right', aspect.ratio = 1,
    legend.title = element_blank(),
    panel.border = element_rect(colour = "black", fill=NA, size=1)) +
  labs(x = "Wavelength (nm)", y = "Mean Residue Ellipticity * 10^-3")

Raw

setwd("C:/Users/Vincent/OneDrive/Documents/Boothby Lab Work/Motif Synergy Project/CD Spec")

file <- read_excel('BeStSel Deconv.xlsx')
file <- subset(file, Protein == "Ta-11mer")

file <- subset(file, Cosolute != 'Urea')
file$Cosolute <- factor(file$Cosolute, ordered = T, levels = c("Buffer", "Trehalose", "Sucrose", "Betaine"))

Decon <- ggplot(data = file) +
  geom_col(position = 'fill', color = 'black', aes(x = Cosolute, y = Value, fill = Structure)) +
  scale_fill_manual(values = c("#e9e9e9", "#3c6e71", "#aeaeae", "#353535", "#284b63")) +
  theme_bw() +
  theme(
    plot.title = element_text(hjust = 0.05, size = 14, vjust = -8),
    axis.title = element_text(size =14),
    axis.text.y = element_text(size =10),
    axis.text.x = element_text(size = 10),
    legend.text=element_text(size=14),
    legend.position = 'right', aspect.ratio = 1.2,
    legend.title = element_blank(),
    panel.border = element_rect(colour = "black", fill=NA, size=1)) +
  labs(y = "Fractional Structure")

Decon

Raw | Decon
```
