## Supplementary material for "LEA_4 motifs function alone and in conjunction with synergistic cosolutes to protect a labile enzyme during desiccation": File S2: LEA_4 TFE.py

```
def aminoToEnergy(amino, cosolute):
    # Note: NO values given for Cys. Set as 0
    # Determines which cosolute to calculate from literature values

    treDict = {
        "A": 59.3,  "F": 57.1,  "L": 190.8, "I": 221.3, "V": 153.0,
        "P": 135.7, "M": 246.56, "G": 0, "S": -1.58, "T": 13.20,
        "Q": -23.98, "N": -123.1, "D": -67.6, "E": -86.7, "H": -68.0,
        "K": 204.3, "R": 193.9, "C": 0, "W": 35.9, "Y": -96.1
    }

    tmaoDict = {
        "A": -14.64, "F": -9.32, "L": 11.62, "I": -25.43, "V": -1.02,
        "P": -137.73, "M": -7.65, "W": -152.87, "G": 0, "S": -39.05,
        "T": 3.57, "Y": -114.32, "Q": 41.41, "N": 55.69, "D": -66.67,
        "E": -83.25, "H": 42.07, "K": -110.23, "R": -109.27, "C": 0,
    }

    sarDict = {
        "A": 10.91, "F": -12.64, "L": 38.33, "I": 39.98, "V": 29.32,
        "P": -32.23, "M": 8.18, "W": -113.03, "G": 0, "S": -27.98,
        "T": -7.54, "Y": -26.37, "Q": -10.19, "N": -40.93, "D": -14.20,
        "E": -12.61, "H": -20.80, "K": -27.42, "R": -32.24, "C": 0,
    }

    betDict = {
        "A": 4.77, "F": -112.93, "L": -17.73, "I": -1.27, "V": -19.63,
        "P": -125.16, "M": -14.16, "W": -369.93, "G": 0, "S": -241.85,
        "T": 0.33, "Y": -213.09, "Q": 7.57, "N": 33.17, "D": -116.56,
        "E": -112.08, "H": -35.97, "K": -171.99, "R": -109.45, "C": 0
    }

    sorDict = {
        "A": 16.57, "F": 26.38, "L": 39.07, "I": 36.90, "V": 24.65,
        "P": -4.48, "M": 20.97, "W": -67.23, "G": 0, "S": -1.58,
        "T": 13.20, "Y": -53.50, "Q": -23.98, "N": -21.21, "D": -83.88,
        "E": -70.05, "H": -42.45, "K": -32.47, "R": -24.65, "C": 0,
    }

    sucDict = {
        "A": 22.05, "F": -96.35, "L": 31.11, "I": 28.12, "V": 33.92,
        "P": -73.02, "M": -6.66, "G": 0, "S": -2.79, "T": 20.82,
        "Q": -40.87, "N": -28.28, "D": -37.17, "E": -41.65, "H": -118.66,
        "K": -39.60, "R": -79.32, "C": 0, "W": -215.27, "Y": -78.41
    }

    ureaDict = {
        "A": -4.69, "F": -83.11, "L": -54.57, "I": -38.43, "V": -21.65,
        "P": -17.65, "M": -48.34, "G": 0, "S": -20.56, "T": 13.20,
        "Q": -54.81, "N": -38.79, "D": 3.55, "E": 0.62, "H": -50.51,
        "K": -22.79, "R": -21.17, "C": 0, "W": -141.46, "Y": -45.08
    }

    proDict = {
        "A": -0.07, "F": -71.26, "L": 4.77, "I": -2.72, "V": 7.86,
        "P": -63.96, "M": -35.12, "G": 0, "S": -33.49, "T": -18.33,
        "Q": -32.36, "N": -17.71, "D": -90.51, "E": -89.17, "H": -45.10,
        "K": -59.87, "R": -60.18, "C": 0, "W": -198.37, "Y": -138.41
    }

    glyDict = {
        "A": 7.77, "F": 59.78, "L": -34.41, "I": 36.23, "V": -1.37,
        "P": -60.54, "M": 13.88, "W": -122.65, "G": 0, "S": 6.31,
        "T": 17.54, "Y": -149.50, "Q": -2.75, "N": 51.57, "D": -85.45,
        "E": -74.20, "H": -17.16, "K": -34, "R": -30.74, "C": 0
    }

    if cosolute == "trehalose":
        return treDict[amino]
    if cosolute == "trehaloseBack":
        return treDict[amino] + 35
    if cosolute == "tmao":
        return tmaoDict[amino]
    if cosolute == "tmaoBack":
        return tmaoDict[amino] + 90
    if cosolute == "sarcosine":
        return sarDict[amino]
    if cosolute == "sarcosineBack":
        return sarDict[amino] + 52
    if cosolute == "betaine":
        return betDict[amino]
    if cosolute == "betaineBack":
        return betDict[amino] + 67
    if cosolute == "sorbitol":
        return sorDict[amino]
    if cosolute == "sorbitolBack":
        return sorDict[amino] + 35
    if cosolute == "sucrose":
        return sucDict[amino]
    if cosolute == "sucroseBack":
        return sucDict[amino] + 62
    if cosolute == "urea":
        return ureaDict[amino]
    if cosolute == "ureaBack":
        return ureaDict[amino] - 39
    if cosolute == "proline":
        return proDict[amino]
    if cosolute == "prolineBack":
        return proDict[amino] + 48
    if cosolute == "glycerol":
        return glyDict[amino]
    if cosolute == "glycerolBack":
        return glyDict[amino] + 14
    
cosoluteList = ["trehalose", "sucrose", "betaine"]
sequence = "AKDKVGDAYDA"

for cosolute in cosoluteList:
    sum = 0
    for aa in sequence:
        sum = sum + aminoToEnergy(aa, cosolute + "Back")
    print(cosolute + ": " + str(sum))
```
