## Supplementary material for "LEA_4 motifs function alone and in conjunction with synergistic cosolutes to protect a labile enzyme during desiccation": File S2: TFE Combined Stripchart.R

```
library(readr)
library(dplyr)
library(ggplot2)
library(readxl)
library(ggpmisc)

setwd("C:/Users/Vincent/OneDrive/Documents/Boothby Lab Work/Motif Synergy Project/")

file <- read_excel('Data Master.xlsx')

file <- subset(file, Solute == "Trehalose" | Solute == "Sucrose" | Solute == "Betaine")
file <- subset(file, Peptide != "Ce-22mer" & Peptide != "Ce-44mer" & Peptide != "At-44mer" & Peptide != "At-22mer")

file$Peptide <- factor(file$Peptide, levels = c("Af-11mer", "At-11mer", "Ce-11mer", "Hb-11mer", "Rt-11mer", "Ta-11mer", "Tg-11mer"), ordered = T)

Solutes <- unique(file$Solute)
Peptides <- c("Af-11mer", "At-11mer", "Ce-11mer", "Hb-11mer", "Rt-11mer", "Ta-11mer", "Tg-11mer")

Lines <- c(1.5, 2.5, 3.5, 4.5, 5.5, 6.5, 7.5, 8.5, 9.5, 10.5)

stripchart(`ΔG` ~ Peptide, 
           data = file, 
           pch = 21,
           group.names = Peptides,
           method = "jitter",
           col = "black",
           cex = 2,
           bg = rep(c("#C8553D", "#F28F3B", "#93B7BE")),
           ylim = c(-400, 400),
           ylab = "Change in MRE at 222/208nm",
           vertical = T,
           axes = F)
abline(v = Lines)
abline(h = 0, lty = 5)
axis(2)
box(lwd = 2)

ggplot(data = file) +
  geom_point(position = position_dodge(0.75), aes(x = Peptide, y = ΔG, fill = Solute), size = 3, pch = 21, color = "black") +
  geom_hline(yintercept=0, linetype = "dashed") +
  geom_vline(xintercept=c(1.5, 2.5, 3.5, 4.5, 5.5, 6.5)) +
  scale_color_manual(values = c("#93B7BE", "#F28F3B", "#C8553D")) +
  scale_fill_manual(values = c("#93B7BE", "#F28F3B", "#C8553D")) +
  theme_bw() +
  theme(
    plot.title = element_text(hjust = 0.05, size = 14, vjust = -8),
    axis.title = element_text(size =14),
    axis.text.y = element_text(size =10),
    axis.text.x = element_text(size = 10),
    legend.text=element_text(size=14),
    legend.position = "none",
    aspect.ratio = 0.6,
    legend.title = element_blank(),
    panel.border = element_rect(colour = "black", fill=NA, size=1),
    panel.grid.major = element_blank(), 
    panel.grid.minor = element_blank()) +
  labs(y = "Changes in Rg (Å)")
```
