## Supplementary material for "LEA_4 motifs function alone and in conjunction with synergistic cosolutes to protect a labile enzyme during desiccation": File S2: TFE Synergy Barplot.R

```
library(tidyverse)
library(readxl)
library(ggpmisc)
library(patchwork)

names <- c("Trehalose", "Sucrose", "Betaine")

setwd("C:/Users/Vincent/OneDrive/Documents/Boothby Lab Work/Motif Synergy Project")
file <- read_excel("Data Master.xlsx")
file <- subset(file, Peptide == "Af-11mer" | Peptide == "At-11mer" | Peptide == "Ce-11mer" | Peptide == "Hb-11mer" | Peptide == "Rt-11mer" | Peptide == "Ta-11mer" | Peptide == "Tg-11mer")

Synergy <- c(mean(subset(file, Solute == "Trehalose")$Synergy),
             mean(subset(file, Solute == "Sucrose")$Synergy),
             mean(subset(file, Solute == "Betaine")$Synergy))  

Error <- c(sd(subset(file, Solute == "Trehalose")$Synergy),
           sd(subset(file, Solute == "Sucrose")$Synergy),
           sd(subset(file, Solute == "Betaine")$Synergy))

temp <- data.frame(names, Synergy, Error)
cols <- c("#93B7BE", "#F28F3B", "#C8553D")

Syn <- ggplot(data = temp) +
  geom_bar(position = position_dodge(), stat = 'identity', color = 'black', aes(x = ordered(names, levels = names), y = `Synergy`, fill = names)) +
  theme_bw()+
  theme(
    plot.title = element_text(hjust = 0.05, size = 14, vjust = -8),
    axis.title = element_text(size =14),
    axis.text.y = element_text(size =10),
    axis.text.x = element_text(size = 10),
    legend.text=element_text(size=14),
    legend.position = 'none', aspect.ratio = 1,
    legend.title = element_blank(),
    panel.border = element_rect(colour = "black", fill=NA, size=1)) +
  geom_errorbar(aes(x = `names`, ymin = `Synergy` - `Error`, ymax = `Synergy` + `Error`), width = 0.2, size = 0.8) +
  scale_fill_manual(values = cols) +
  labs(x = '', y = 'Mean Synergy')
  
Syn

Protection <- c(mean(subset(file, Solute == "Trehalose")$Protection),
             mean(subset(file, Solute == "Sucrose")$Protection),
             mean(subset(file, Solute == "Betaine")$Protection))  

Error <- c(sd(subset(file, Solute == "Trehalose")$Protection),
           sd(subset(file, Solute == "Sucrose")$Protection),
           sd(subset(file, Solute == "Betaine")$Protection))

temp <- data.frame(names, Protection, Error)
cols <- c("#93B7BE", "#F28F3B", "#C8553D")

Pro <- ggplot(data = temp) +
  geom_bar(position = position_dodge(), stat = 'identity', color = 'black', aes(x = ordered(names, levels = names), y = `Protection`, fill = names)) +
  theme_bw()+
  theme(
    plot.title = element_text(hjust = 0.05, size = 14, vjust = -8),
    axis.title = element_text(size =14),
    axis.text.y = element_text(size =10),
    axis.text.x = element_text(size = 10),
    legend.text=element_text(size=14),
    legend.position = 'none', aspect.ratio = 1,
    legend.title = element_blank(),
    panel.border = element_rect(colour = "black", fill=NA, size=1)) +
  geom_errorbar(aes(x = `names`, ymin = `Protection` - `Error`, ymax = `Protection` + `Error`), width = 0.2, size = 0.8) +
  scale_fill_manual(values = cols) +
  labs(x = '', y = 'Mean Protection')

Pro

Syn | Pro
```
