## Supplementary material for "LEA_4 motifs function alone and in conjunction with synergistic cosolutes to protect a labile enzyme during desiccation": File S2: Cider Correlations.R

```
library(tidyverse)
library(readxl)
library(ggpmisc)

setwd("C:/Users/Vincent/OneDrive/Documents/Boothby Lab Work/Motif Synergy Project/Informatics")

file <- read_excel('CIDER.xlsx')

# ttmers <- subset(file, Peptide == "At-22mer" | Peptide == "Ce-22mer")
# ffmers <- subset(file, Peptide == "At-44mer" | Peptide == "Ce-44mer")

file$Active <- file$'Trehalose'

cor <- cor.test(file$Interaction, file$Active, method = "pearson")
r <- unname(cor$estimate)
r^2
cor$p.value

text <- c(substr(paste(expression(R^2), "=", r^2), 1, 12), substr(paste("p", "=", cor$p.value), 1, 10))
ypos <- c(0, 0.05)

test <- data.frame(text, ypos)

annotations <- data.frame(xpos = c(Inf, Inf),
ypos =  c(Inf, Inf),
annotateText = c(substr(paste(expression(R^2), "=", r^2), 1, 12), substr(paste("p", "=", cor$p.value), 1, 10)),
hjustvar = c(1.05, 1.05) ,
vjustvar = c(1.5, 2.9))

ggplot() +
  geom_point(data = file, aes(x = Interaction, y = Active, fill = homo), size = 3.5, color = "black", shape = 21) +
  geom_smooth(data = file, method = "lm", aes(x = Interaction, y = Active), se = F, color = "black") +
  geom_text(data=annotations,aes(x=xpos,y=ypos,hjust=hjustvar,vjust=vjustvar,label=annotateText)) +
  theme_bw()+
  theme(
    plot.title = element_text(hjust = 0.05, size = 14, vjust = -1.5),
    axis.title = element_text(size =10),
    axis.text.y = element_text(size =10),
    axis.text.x = element_text(size = 10),
    legend.text=element_text(size=14),
    legend.position = 'right', aspect.ratio = 1,
    legend.title = element_text(size = 10),
    panel.border = element_rect(colour = "black", fill=NA, size=1))+
  scale_fill_gradient2(low = "darkorchid4", high = "darkgreen", lim = c(-2.5, 2.5)) +
  labs(x = "Motif:CS ε", y = "Fold Synergy", fill = expression(Homo[ε]), title = "Trehalose")
```
