## Supplementary material for "LEA_4 motifs function alone and in conjunction with synergistic cosolutes to protect a labile enzyme during desiccation": File S2: CIDER Visualization Plot.R

```
library(tidyverse)
library(readxl)
library(ggpmisc)

setwd("C:/Users/Vincent/OneDrive/Documents/Boothby Lab Work/Motif Synergy Project/Informatics")

file <- read_excel('CIDER.xlsx')

ggplot(data = file) +
  geom_col(position = position_dodge2(width = 0.9), aes(x = Protein, y = κ), color = "black", fill = "grey") +
  theme_bw() +
  theme(
    plot.title = element_text(hjust = 0.05, size = 14, vjust = -8),
    axis.title = element_text(size =14),
    axis.text.y = element_text(size =10),
    axis.text.x = element_text(size = 10),
    legend.text=element_text(size=14),
    legend.position = 'right', aspect.ratio = 0.3,
    legend.title = element_blank(),
    panel.border = element_rect(colour = "black", fill=NA, size=1)) +
  labs(x = "Peptide", y = "κ")

y <- file$`Hydropathy`
x <- file$Protein

stripchart(y ~ x, 
           pch = 21,
           col = "black",
           cex = 1.5,
           bg = "#284b63",
           ylim = c(2, 4.5),
           ylab = "Kyte-Doolittle Hydropathy",
           vertical = T,
           axes = F)
abline(h = 4.5, lty = 5)
axis(2)
box(lwd = 2)

y <- file$`κ`
x <- file$Protein

stripchart(y ~ x, 
           pch = 21,
           col = "black",
           cex = 1.5,
           bg = "#284b63",
           ylim = c(0, 1),
           ylab = "κ Value",
           vertical = T,
           axes = F)
axis(2)
box(lwd = 2)
```
