## Supplementary material for "LEA_4 motifs function alone and in conjunction with synergistic cosolutes to protect a labile enzyme during desiccation": File S2: Kratky.R

```
library(readr)
library(dplyr)
library(ggplot2)
library(readxl)
library(ggpmisc)
library(pander)
library(patchwork)

setwd("C:/Users/Vincent/OneDrive/Documents/Boothby Lab Work/Motif Synergy Project/SAXS/csv")

list <- c("Af-11", "At-11", "Ce-11", "Hb-11", "Rt-11", "Ta-11", "Tg-11")

Kratky_list <- c()
Guinier_list <- c()
for(i in 1:length(list))
{
  name <- paste(list[i], ".csv", sep = "")
  name_guinier <- paste(list[i], " Guinier", ".csv", sep ="")
  file <- read.csv(name)
  file_guinier <- read.csv(name_guinier)
  file$Kratky <- (file$I.Q. * file$Q * file$Q)
  
  krat <- ggplot(data = file) +
    geom_point(aes(x = Q, y = Kratky), pch = 21, color = "black", bg = "white") +
    theme_bw()+
    theme(
      plot.title = element_text(size =8, v = -8, h = 0.05),
      axis.title = element_text(size =8),
      axis.text.y = element_text(size =8),
      axis.text.x = element_text(size = 8),
      legend.text = element_blank(),
      legend.position = 'right', aspect.ratio = 1,
      legend.title = element_blank(),
      panel.border = element_rect(colour = "black", fill=NA, size=1)) +
    ylim(0, 0.04) +
    xlim(0, 0.45) +
    labs(x = expression('q' * ' ' * (Å ^ -1)), y = expression('I(q)' * (q ^ 2)), title = list[i])
  
  guin <- ggplot() +
    geom_point(data = file, aes(x = Q * Q, y = log(I.Q.)), pch = 21, color = "black", bg = "white") +
    geom_line(data = file_guinier, aes(x = `q..2_fit`, y = `ln.I.q.._fit`), color = "red", size = 1) +
    theme_bw()+
    theme(
      plot.title = element_blank(),
      axis.title = element_text(size =8),
      axis.text.y = element_text(size =8),
      axis.text.x = element_text(size = 8),
      legend.text = element_blank(),
      legend.position = 'right', aspect.ratio = 0.5,
      legend.title = element_blank(),
      panel.border = element_rect(colour = "black", fill=NA, size=1)) +
    labs(x = expression(q^2 * ' ' * (Å ^ -2)), y = 'ln(I(q))') +
    xlim(0, 0.02) +
    ylim(-2, 0.5)
  
  Kratky_list[[i]] <- krat
  Guinier_list[[i]] <- guin
}

(Kratky_list[[1]] | Kratky_list[[2]] | Kratky_list[[3]]) / (Kratky_list[[4]] | Kratky_list[[5]] | Kratky_list[[6]]) / (Kratky_list[[7]] | Kratky_list[[7]] | Kratky_list[[7]])
(Guinier_list[[1]] | Guinier_list[[2]] | Guinier_list[[3]]) / (Guinier_list[[4]] | Guinier_list[[5]] | Guinier_list[[6]]) / (Guinier_list[[7]] | Guinier_list[[7]] | Guinier_list[[7]])

((Kratky_list[[1]]/Guinier_list[[1]]) | (Kratky_list[[2]]/Guinier_list[[2]]) | (Kratky_list[[3]]/Guinier_list[[3]])) / ((Kratky_list[[4]]/Guinier_list[[4]]) | (Kratky_list[[5]]/Guinier_list[[5]]) | (Kratky_list[[6]]/Guinier_list[[6]])) / ((Kratky_list[[7]]/Guinier_list[[7]]) | (Kratky_list[[7]]/Guinier_list[[7]]) | (Kratky_list[[7]]/Guinier_list[[7]]))
  
```
