## Supplementary material for "LEA_4 motifs function alone and in conjunction with synergistic cosolutes to protect a labile enzyme during desiccation": File S2: Mixing.R

```
library(readr)
library(dplyr)
library(ggplot2)
library(readxl)
library(ggpmisc)

setwd("C:/Users/Vincent/OneDrive/Documents/Boothby Lab Work/Motif Synergy Project/CS Assay")

file <- read_excel('Mixing Data Compiled.xlsx', sheet = 'Tg-11mer')

file$Solute = factor(file$Solute, levels = c("Trehalose", "Sucrose", "Formate", "Spermidine", "y-Glutamylalanine", "No-Acetylornithine", "Betaine"))

file$Type = factor(file$Type, levels = c("Solute", "Peptide", "Expected", "Actual"))

New <- ggplot(data = file, aes(x = Solute, y = Result, fill = Type)) +
  geom_col(data = file %>% filter(Type == 'Solute'| Type == 'Actual'), 
           position = position_dodge(width = .9), 
           color = 'black')+
  geom_col(data = file %>% filter(Type == 'Expected'| Type == 'Actual'), 
           position = position_dodge(width = .9), 
           alpha = .6, 
           color = 'black')+
  labs(title = "Tg-11mer", y = "Protection (%)") +
  scale_y_continuous(name = '%Protection',
                     breaks = seq(0,120,20),
                     limits = c(0,120)) +
  scale_fill_manual(values = c("darkgrey", "white", "grey")) +
  theme_bw() +
  theme(
    plot.title = element_text(hjust = 0.05, size = 14, vjust = -8),
    axis.title = element_text(size =14),
    axis.text.y = element_text(size =10),
    axis.text.x = element_text(size = 10),
    legend.text=element_text(size=14),
    legend.position = 'right', aspect.ratio = 0.5,
    legend.title = element_blank(),
    panel.border = element_rect(colour = "black", fill=NA, size=1)) +
    geom_errorbar(data = file %>% filter(Type == 'Expected'| Type == 'Actual'),
          aes(ymin = Result - Error, ymax = Result + Error), width = .2, position = position_dodge(.9))

New

file <- read_excel('Example.xlsx')

file$Solute = factor(file$Solute, levels = c("Additive", "Synergy", "Antagonism"))

file$Type = factor(file$Type, levels = c("Solute", "Peptide", "Expected", "Actual"))

New <- ggplot(data = file, aes(x = Solute, y = Result, fill = Type)) +
  geom_col(data = file %>% filter(Type == 'Solute'| Type == 'Actual'), 
           position = position_dodge(width = .9), 
           color = 'black')+
  geom_col(data = file %>% filter(Type == 'Expected'| Type == 'Actual'), 
           position = position_dodge(width = .9), 
           alpha = .6, 
           color = 'black')+
  labs(title = "Example", y = "Protection (%)") +
  scale_y_continuous(name = '%Protection',
                     breaks = seq(0,120,20),
                     limits = c(0,120)) +
  scale_fill_manual(values = c("darkgrey", "white", "grey")) +
  theme_bw() +
  theme(
    plot.title = element_text(hjust = 0.05, size = 14, vjust = -8),
    axis.title = element_text(size =14),
    axis.text.y = element_text(size =10),
    axis.text.x = element_text(size = 10),
    legend.text=element_text(size=14),
    legend.position = 'right', aspect.ratio = 0.5,
    legend.title = element_blank(),
    panel.border = element_rect(colour = "black", fill=NA, size=1)) +
  geom_errorbar(data = file %>% filter(Type == 'Expected'| Type == 'Actual'),
                aes(ymin = Result - Error, ymax = Result + Error), width = .2, position = position_dodge(.9))

New
```
