## Supplementary material for "LEA_4 motifs function alone and in conjunction with synergistic cosolutes to protect a labile enzyme during desiccation": File S2: Dichro Patchwork.R

```
library(tidyverse)
library(readxl)
library(ggpmisc)
library(patchwork)

setwd("C:/Users/Vincent/OneDrive/Documents/Boothby Lab Work/Motif Synergy Project/CD Spec")

file <- read_excel('Dichroweb Deconv.xlsx', sheet = 'Sheet3')
file <- subset(file, Cosolute != 'Urea')

proteins <- c("Af-11mer", "At-11mer", "Ce-11mer", "Hb-11mer", "Rt-11mer", "Ta-11mer", "Tg-11mer")
Decon_list <- c()

for(i in 1:length(proteins))
{
  temp <- subset(file, Protein == proteins[i])
  
  temp$Cosolute <- factor(temp$Cosolute, ordered = T, levels = c("Buffer", "Trehalose", "Sucrose", "Betaine"))

  Decon <- ggplot(data = temp) +
    geom_col(position = 'fill', color = 'black', aes(x = Cosolute, y = Value, fill = Structure)) +
    scale_fill_manual(values = c("#448B74", "#A63A2E", "#36628B", "#353535")) +
    theme_bw() +
    theme(
      plot.title = element_text(hjust = 0.05, size = 14, vjust = -8),
      axis.title = element_text(size =8),
      axis.text.y = element_text(size =8),
      axis.text.x = element_text(size = 8),
      legend.text=element_text(size=8),
      legend.position = '', aspect.ratio = 1.2,
      legend.title = element_blank(),
      panel.border = element_rect(colour = "black", fill=NA, size=1)) +
    labs(y = "Fractional Structure")  
  
  Decon_list[[i]] <- Decon
}

(Decon_list[[1]] | Decon_list[[2]] | Decon_list[[3]]) / (Decon_list[[4]] | Decon_list[[5]] | Decon_list[[6]]) / (Decon_list[[7]] | Decon_list[[7]] | Decon_list[[7]])
```
